# Binocular processing facilitates escape behavior through multiple pathways to the superior colliculus

**DOI:** 10.1101/2024.07.22.604589

**Authors:** Robin Broersen, Genevieve Thompson, Felix Thomas, Greg J. Stuart

**Affiliations:** Eccles Institute of Neuroscience, John Curtin School of Medical Research, Australian National University, Canberra, Australian Capital Territory, Australia; Department of Neuroscience, Erasmus MC, Rotterdam, the Netherlands; Department of Physiology, Monash University, Melbourne, Victoria, Australia; Bruce Lefroy Centre for Genetic Health Research, Murdoch Children’s Research Institute, Melbourne, Victoria, Australia

## Abstract

The superior colliculus (SC) is the main brain region regulating innate defensive behaviors to visual threat. Yet, how the SC integrates binocular visual information and to what extent binocular vision drives defensive behaviors is unknown. Here, we show that binocular vision facilitates visually-evoked escape behavior. Furthermore, we find that SC neurons respond to binocular visual input with diverse synaptic and spiking responses, and summate visual inputs largely sublinearly. Using pathway-specific optogenetic silencing we find that contralateral and ipsilateral visual information is carried to binocular SC neurons through retinal, interhemispheric and corticotectal pathways. These pathways carry binocular visual input to the SC in a layer-specific manner, with superficial layers receiving visual information through retinal input, whereas intermediate and deep layers rely on interhemispheric and corticotectal pathways. Together, our data shed light on the cellular and circuit mechanisms underlying binocular visual processing in the SC and its role in escape behavior.

## Introduction

One of the main functions of the brain is to rapidly detect danger and orchestrate appropriate behavioral responses to avoid injury or death. Vision acts as an important sensory modality for detecting danger in many species. In rodents, vision enables the detection of aerial predatory threats, leading to defensive behaviors to avoid detection (freezing) or to flee (escape)^1,2^. An important brain area responsible for controlling these innate defensive behaviors is the superior colliculus (SC)^3–5^, a layered multisensory midbrain structure^6^ also involved in orienting body and eye movements^7–9^, predatory hunting^10–12^, visual spatial attention and perceptual decision-making^13–15^. While neurons located in the superficial layers of the SC exclusively process visual information^16–18^, those in intermediate and deep layers also process auditory and somatosensory information^19,20^.

Visual input is conveyed to the SC by both retinal ganglion cells (RGC)^21,22^ and via input from the primary visual cortex (V1)^23,24^. Following visual processing, the SC regulates the activity in downstream brain areas involved in generating defensive behaviors, which include the lateral posterior thalamic nucleus (LP), the parabigeminal nucleus (PBG) and the periaqueductal gray (PAG)^3,25,26^. Optogenetic activation of the LP or PBG results in freeze or escape responses, respectively^3,4^. Arrest and freezing responses can also be elicited by direct activation of V1 inputs to the SC^27,28^. Furthermore, certain types of SC-projecting RGCs have been shown to regulate innate defensive responses^29^, indicating that visually-evoked activity in the SC is critical for generating defensive responses to visual threat.

Despite a growing interest in the functional role of the SC, a full understanding of how visual input from both eyes is integrated at the level of individual SC neurons is lacking. Binocular vision is functionally relevant in mice and has been shown to be required for successful prey capture^30,31^. This raises the question whether defensive behaviors to visual threats in mice also rely on binocular vision. Binocular neurons, which integrate visual signals from the two eyes, have been found in mouse V1^32,33^ and more recently in SC^34,35^. However, it is unclear how binocular inputs are integrated by SC neurons at the synaptic level and which neural pathways are involved. Furthermore, it is not known whether binocular vision is required for defensive behaviors to visual threats.

To address these questions, here we combine *in vivo* and *in vitro* electrophysiology with optogenetics and behavior to investigate binocular integration in the mouse SC and its role in defensive responses to visual threat. We find that binocular vision is required for defensive responses to visual threat, facilitating escape responses. At the cellular level, almost half of the neurons in the region of the SC receiving visual input from the binocular part of the visual field were binocular. Synaptic and spiking responses to stimulation of the eyes in binocular neurons were diverse, containing both excitatory and inhibitory components, with integration of binocular responses found to be sublinear. Finally, we show that contralateral and ipsilateral visual input is conveyed to the SC by retinal, interhemispheric commissural and corticotectal projections in a layer-specific manner. Together, our results shed light on the mechanistic principles underlying binocular processing in the SC and highlight its role in defensive responses to visual threats.

## Results

### Binocular vision aids detection of visual threat

To investigate behavioral responses to visual threat, mice were placed in an open field enclosure and presented with a rapidly expanding (“looming”) black spot projected from above on a gray background, representing a rapidly approaching aerial predator (Figure 1A). In response to this stimulus, control (Ctrl) mice escaped to a shelter on the majority of trials (Figure 1B), consistent with previous work^1,2^. In contrast, mice that had undergone monocular enucleation (Enuc), and therefore lacked binocular vision, showed reduced escape responses and increased freezing (Figure 1B-D, Video S1). Enucleation did not impact on the ability of mice to detect the visual stimulus, as the percentage of trials where no response was observed was similar between control and enucleated mice (Ctrl: 3% vs. Enuc: 2%). During escape responses, enucleated mice exhibited a reduced peak velocity and a longer latency to escape, although the length of escape trajectory was unchanged (Figure 1E-I). Differences between control and enucleated mice were also observed during exposure to sweeping spots that typically lead to freezing^1^ (Figure S1). These data show that defensive responses to acute visual threat depend on binocular vision.

**Figure 1.**
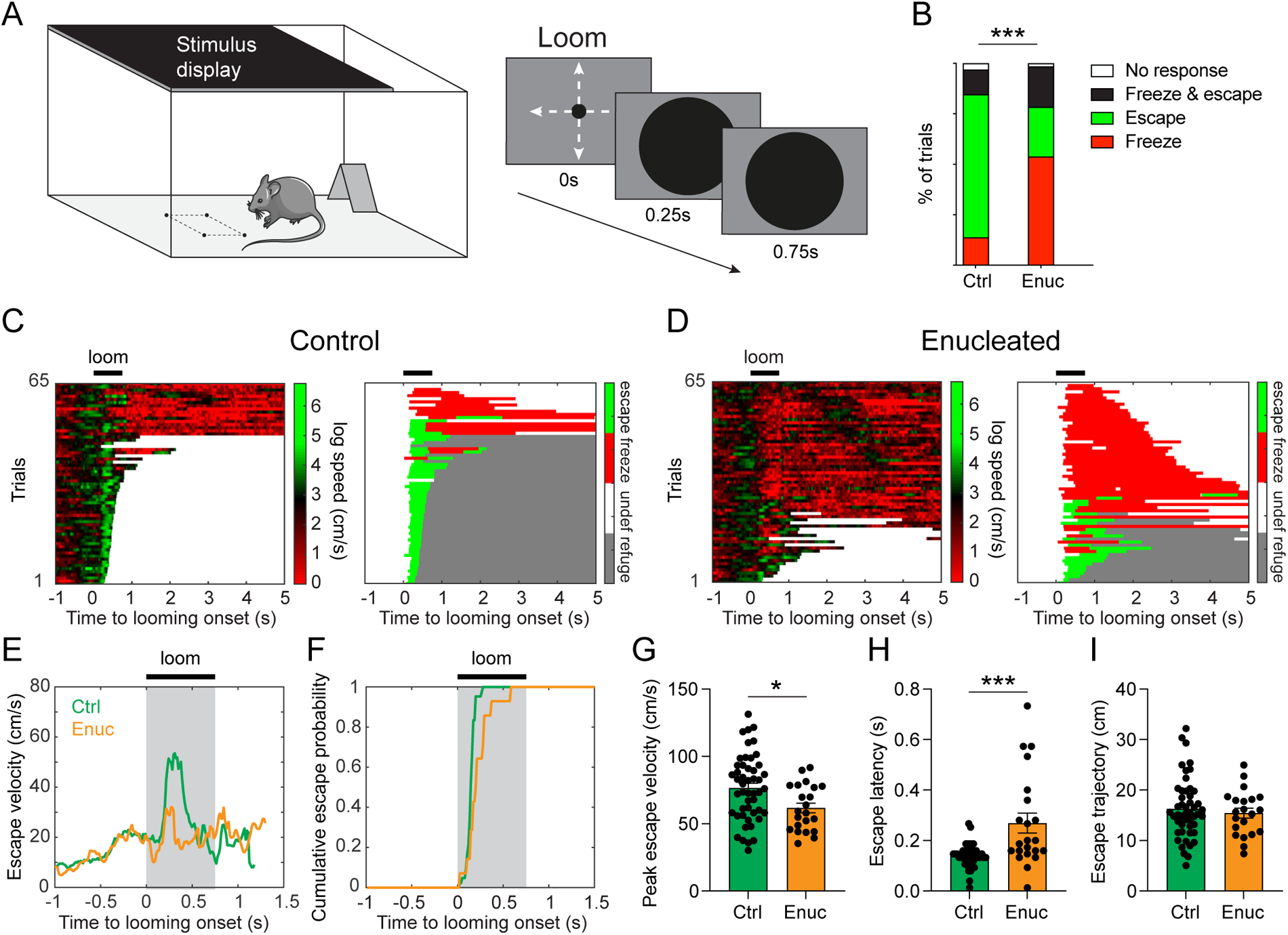
Binocular vision facilitates defensive responses to a looming spot. (A) Experimental setup for measuring defensive responses to visual stimuli. A single overhead looming spot, which expands (0-250ms) and remains stationary (250-750ms), was started when mice enter the center of the arena, indicated by four black dots on the floor. (B) Behavioral responses of control (Ctrl) and enucleated (Enuc) mice to looming stimuli across all trials are different (*χ*^2^(3) = 31.4, p < 0.001, chi-square test, n=65 trials from 15 Ctrl mice, n=65 trials from 17 Enuc mice). (C) Left: Heat map visualizing control mouse movement speed relative to looming onset across all trials. Right: Classification of responses relative to looming onset. Undef (undefined) indicates time periods where behavior was not defined. (D) Same as in C, but for enucleated mice. (E) Median velocity on escape trials relative to looming onset. (F) Cumulative escape probability relative to looming onset. (G-I) Peak escape velocity (G; *t*(69) = 2.595, p = 0.012, unpaired *t*-test, two-tailed, Ctrl: n=49 trials; Enuc: n=22 trials), escape latency (H; U = 226.5, p < 0.001, Mann-Whitney *U* test, two-tailed, Ctrl: n=48 trials; Enuc: n=22 trials) and escape trajectory (*t*(68) = 0.578, p = 0.565, unpaired *t*-test, two-tailed, Ctrl: n=49 trials; Enuc: n=21 trials) on escape trials. Data points show individual trials. Bar plots bars show mean ± SEM. *p < 0.05, **p < 0.01, ***p < 0.001.

### Synaptic processing of binocular vision in the SC

These findings led us to ask how binocular visual inputs are processed in the SC. To investigate binocular processing at the synaptic level, we performed *in vivo* whole-cell recordings from the anteromedial region of the SC which processes information from the binocular part of the visual field in anesthetized mice. To investigate synaptic responses evoked by stimulation of each eye separately or together we used brief (20 ms) light flashes from light-emitting diodes (LEDs) placed in front of the eyes (Figure 2A). Receiver operating characteristic analysis revealed that approximately half of recorded neurons (47%; n=21 out of 45 neurons from 27 mice) responded to both ipsilateral and contralateral eye stimulation and hence were binocular. Forty percent (n=18) of neurons responded to contralateral eye stimulation only, and hence were monocular, and 13% (n=6) did not respond to either eye. Active and passive electrophysiological properties of binocular, monocular and non-responsive neurons were similar (Figure S2A-J, Table S1). Given that previous work indicates that there are four main cell types in the superficial SC (wide-field, narrow-field, stellate and horizontal cells)^36,37^, we classified binocular neurons in the superficial SC (0-600 µm from the dorsal surface, n=32) into four groups based on their electrophysiological properties using k-means clustering. Binocular neurons were found in all clusters, suggesting that all four cell types in the superficial SC contain binocular neurons (Figure S2K). Using principal component analysis, we found that neurons that clustered together occupied similar regions in the principal component parameter space, further supporting the idea that they represent different neuronal populations (Figure S2L).

**Figure 2.**
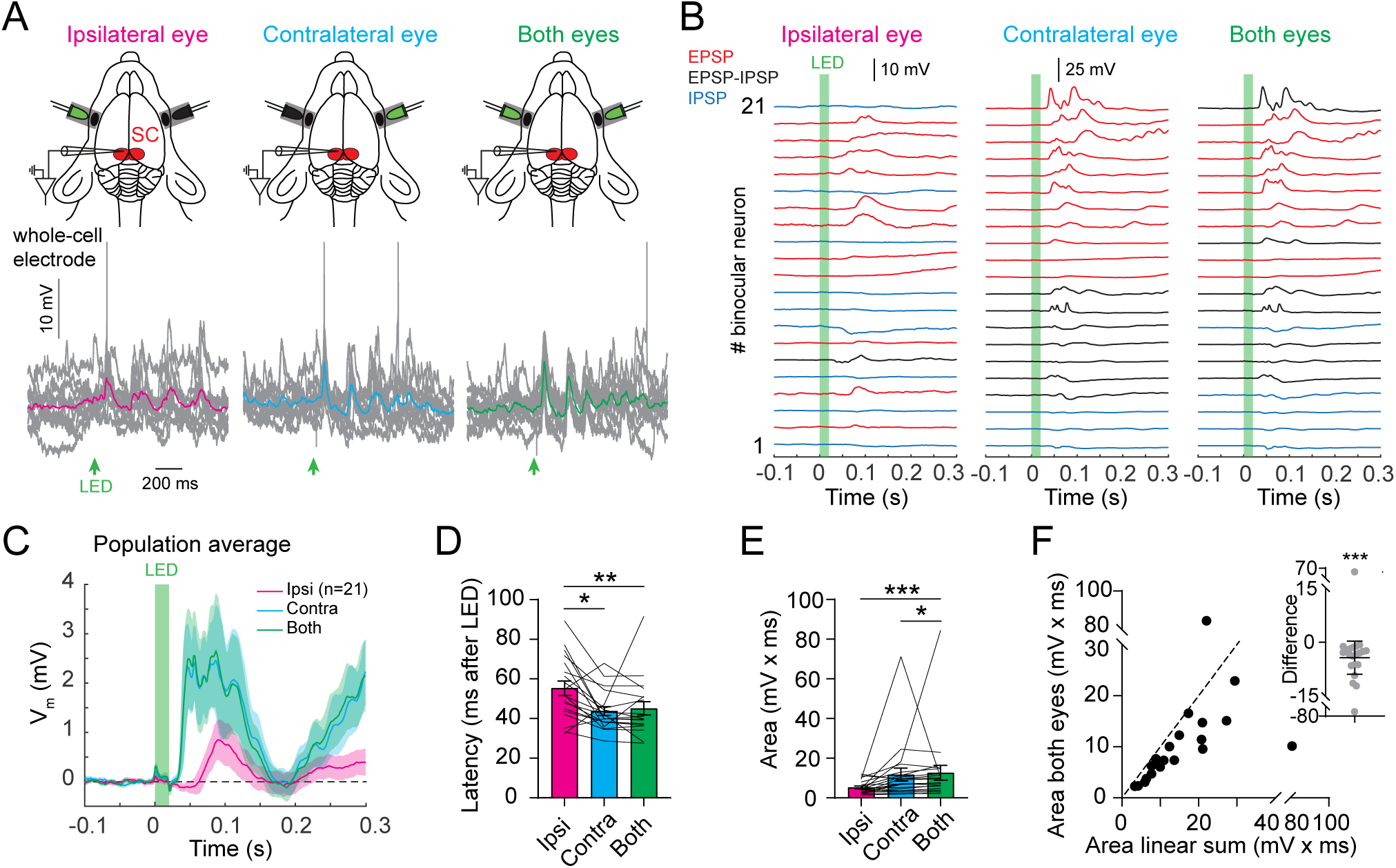
Synaptic processing of binocular visual input in superior colliculus neurons. (A) Top: Schematic of *in vivo* whole-cell recording from the SC (red) during eye-specific stimulation with LED flashes (green). Bottom: Membrane potential of an example neuron during stimulation of the ipsilateral or contralateral eye alone, or both eyes together. Single trials in grey with colored averages superimposed. Action potentials are truncated. (B) Average responses in individual binocular neurons (n=21) during stimulation of the ipsilateral or contralateral eye alone, or both eyes together. Neurons with EPSPs only in red, IPSPs only in blue, EPSPs and IPSPs in black. LED eye stimulation indicated by the green bar. (C) Population average of the synaptic voltage (V_m_; solid lines) of all binocular neurons during stimulation of the ipsilateral (pink) or contralateral eye alone (blue), or both eyes together (green) together with standard error of the mean (light shading). LED eye stimulation indicated by the green bar. (D) Synaptic response latency is longer during ipsilateral eye stimulation (χ^2^(2) = 14, p < 0.001, ipsi vs. both: p = 0.001, ipsi vs. contra: p = 0.018, contra vs. both: p = 0.99, Friedman test with Dunn’s post-hoc test). (E) Area of synaptic responses during stimulation of both eyes is significantly larger than during ipsi or contra eye stimulation alone (χ^2^(2) = 20.1, p < 0.001, ipsi vs. both: p < 0.001, contra vs. both: p = 0.041, ipsi vs. contra: p = 0.104, Friedman test with Dunn’s post-hoc test). (F) Synaptic response during stimulation of both eyes together is significantly smaller than the linear sum of the ipsi and contra eye response alone, indicating sublinear binocular integration (W = -191, p < 0.001, Wilcoxon matched-pairs signed rank test, two-tailed). Data points and lines show individual neurons. Graphs show mean ± SEM. *p < 0.05, **p < 0.01, ***p < 0.001.

Synaptic responses to ipsilateral and contralateral eye stimulation in binocular SC neurons were diverse, containing both excitatory and inhibitory synaptic components (Figure 2B), with the amplitude and latency of synaptic responses scaling with light intensity (Figure S3). Across the population of all binocular neurons (n=21), SC neurons were on average depolarized by visual input during LED stimulation (Figure 2C). Consistent with this, we found that both contralateral and ipsilateral eye stimulation led to excitatory postsynaptic potentials (EPSPs) in the majority of neurons (52%). Multiphasic EPSP and inhibitory postsynaptic potential (IPSP) responses were evoked by both contra- and ipsilateral eye stimulation in 33% and 55% of neurons, respectively, whereas IPSPs alone were observed in a minority of neurons (contra: 14%; ipsi: 42% of neurons). The synaptic response type evoked by stimulation of the ipsi- and contralateral eye were similar in approximately half of SC binocular neuron (52%). These data show that full-retina stimulation recruits both excitatory and inhibitory inputs that lead to diverse synaptic responses in SC neurons during binocular visual input.

Importantly, we found that the response latency to stimulating the ipsilateral eye alone was longer than that to stimulation of the contralateral eye (Figure 2D), suggesting that ipsilateral visual input takes longer to reach the SC than contralateral visual input. Consistent with this, the time at which the binocular synaptic response differed significantly from the contralateral response occurred on average 78 ± 13 ms after LED onset, similar to the latency of the ipsilateral response (*t*(6) = 1.756, p = 0.13, paired *t*-test, two-tailed, n=7 neurons). While the synaptic response to contralateral visual input was greater than that to ipsilateral visual input, the ipsilateral eye still contributed significantly to the binocular response, as the synaptic response area during binocular stimulation was larger than that during stimulation of the contralateral eye alone (Figure 2E). Finally, we investigated synaptic integration of binocular responses during stimulation of both eyes together, finding that binocular responses were smaller than the linear sum of the ipsi- and contralateral eye response alone (Figure 2F). These data indicate that, similar to primary visual cortex (V1)^33,38^, integration of binocular visual input in SC neurons is sublinear.

### Spike responses closely resemble synaptic responses

To investigate how binocular synaptic inputs are translated into spiking we performing juxtasomal recordings from SC neurons during LED flashes (n=77 neurons from 5 mice). Consistent with the distribution of synaptic responses, we found that 49% of neurons were binocular, 29% were monocular (23% contra-responding, 5% ipsi-responding) and 22% did not respond to LED flashes. Consistent with the diversity of synaptic responses in binocular neurons (Figure 2B), we found a diversity of spiking responses in binocular neurons, which could be classified in four types (Figure S4), as observed previously^34^. Forty percent of binocular cells evoked spiking in responses to stimulation of both the ipsi- and contralateral eye (Figure 3A; type 1). Importantly, we found that these binocular neurons were observed across all layers of the SC, up to 1.4 mm from the SC surface (Figure 3B). Also consistent with synaptic responses, we found that type 1 binocular SC neurons spiked more stronger during contralateral compared to ipsilateral eye stimulation (Figure 3C), resulting in the majority of binocular neurons having a positive ocular dominance index (ODI) score (Figure 3D). Finally, as with synaptic responses, we found that spike latency was longer during ipsilateral eye stimulation compared to contralateral eye stimulation (Figure 3E), with integration of ipsilateral and contralateral eye spiking sublinear in most binocular SC cells (Figure 3F).

**Figure 3.**
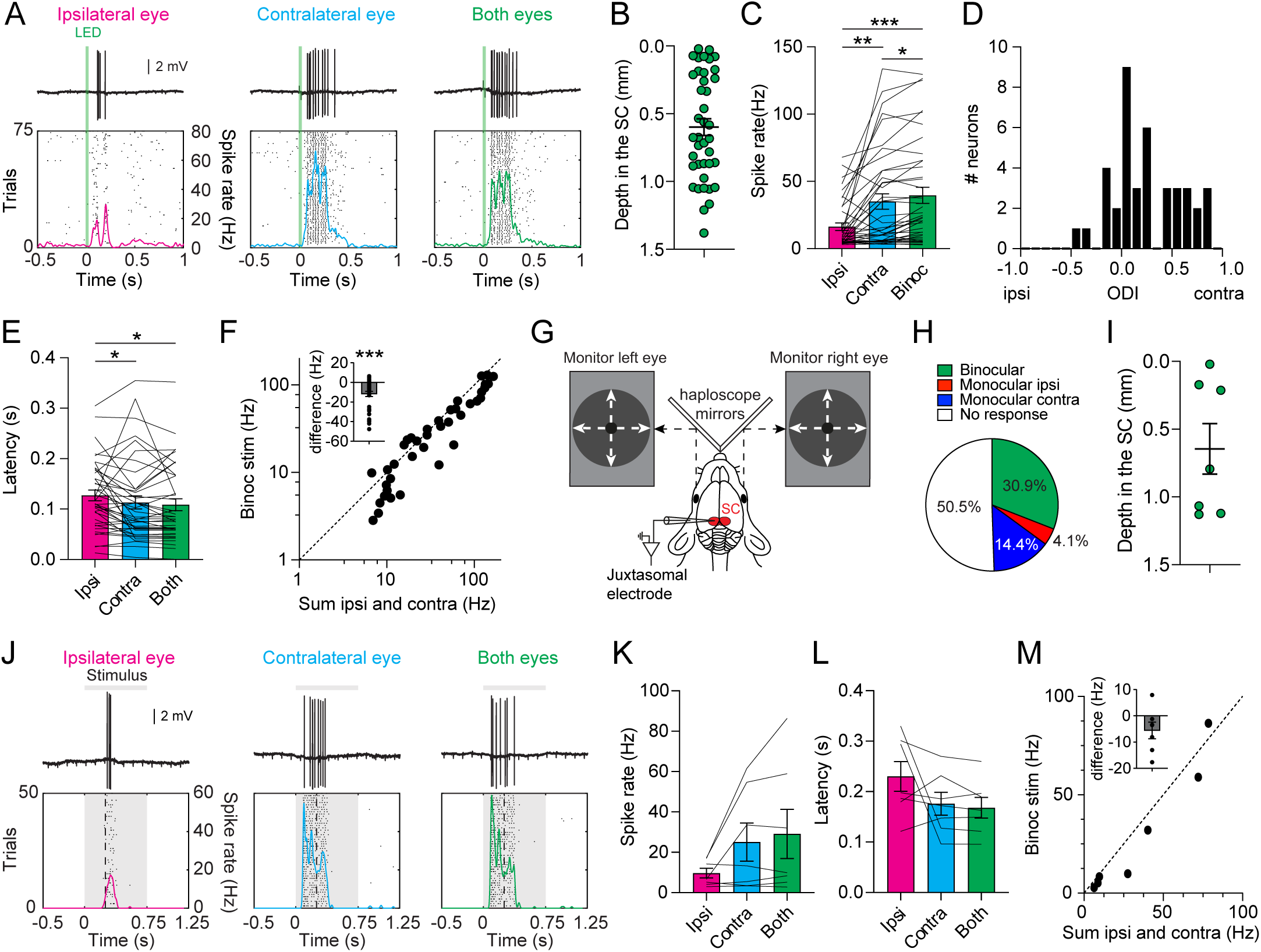
Spike activity during binocular visual inputs in superior colliculus neurons. (A) Top: Spikes in an example binocular neuron during stimulation of the ipsilateral (left, pink) or contralateral (middle, blue) eye alone, or both eyes together (right, green). Bottom: Peristimulus time histograms with each dot representing a spike. Colored lines show spike rate (right Y-axis). (B) Depth of binocular neurons (n=40) in the SC. (C) Spike rate for binocular neurons is highest during binocular stimulation, with contralateral eye stimulation alone generating larger responses than ipsilateral eye stimulation (χ^2^(2) = 32.55, p < 0.001, ipsi vs. both: p < 0.001, contra vs. both: p = 0.022, ipsi vs. contra: p < 0.01, Friedman test with Dunn’s post-hoc test). (D) Ocular dominance index (ODI) for binocular neurons. (E) Spike latency is longer during ipsilateral eye stimulation alone (χ^2^(2) = 9.8, p < 0.01, ipsi vs. both: p = 0.011, contra vs. both: p > 0.05, ipsi vs. contra: p < 0.05, Friedman test with Dunn’s post-hoc test). (F) Spike rate during binocular stimulation is lower than the linear sum of spiking responses during ipsilateral and contralateral stimulation alone (W = -590, p < 0.001, Wilcoxon matched-pairs signed rank test, two-tailed). (G) Schematic of *in vivo* juxtasomal recordings with eye-specific looming spot stimuli presented to each eye individually using haploscope mirrors. (H) Distribution of spike responses during looming stimuli (n=97). (I) Depth of binocular neurons (n=7) in the SC. (J) Top: Spiking in a binocular neuron during ipsilateral, contralateral or binocular eye stimulation. Bottom: Peristimulus time histogram. Each dot represents a spike. Colored lines show spike rate (right Y-axis). (K) Spike rate for binocular neurons during looming stimuli presented to the contralateral, ipsilateral or both eyes simultaneously (F = 3.502, p = 0.053, one-way repeated-measures ANOVA, one-tailed). (L) Spiking latency during looming stimuli presented to the contralateral, ipsilateral or both eyes simultaneously (F = 2.313, p = 0.089, one-way repeated-measures ANOVA, one-tailed). (M) Spike rate during binocular stimulation vs. the linear sum of spiking during ipsilateral and contralateral stimulation alone (W = -20, p = 0.055, Wilcoxon matched-pairs signed rank test, one-tailed). Data points and lines show individual neurons. Graphs show mean ± SEM. *p < 0.05, **p < 0.01, ***p < 0.001.

While full-field LED flashes provide a useful stimulus to characterize synaptic input, and can drive SC-dependent arrest behavior^27^, they do not optimally drive SC neurons^39,40^. We therefore investigated neuronal activity during more physiological looming stimuli used to drive escape behavior, as described above (Figure 1). Looming stimuli were presented to the eyes separately using a haploscope (Figure 3G). The distribution of spiking responses (n=97 neurons) to looming stimuli was similar to that during LED stimuli, although there were fewer binocular neurons and more non-responsive neurons (Figure 3H). Similar to responses to LED flashes, binocular responses to looming stimuli were found across all three layers of the SC (Figure 3I). Binocular responses to looming stimuli also showed a trend towards higher spike rates during contralateral and both eye stimulation (Figure 3J-K), and a trend towards longer spike latencies during ipsilateral eye stimulation (Figure 3L). Finally, we found that most binocular responses showed sublinear integration of ipsilateral and contralateral eye spiking responses during looming stimuli (Figure 3M). These data show that binocular neurons in the SC integrate visual input from the ipsilateral and contralateral eye during looming visual stimuli in a similar way to that observed during LED flashes.

### Anatomical analysis of RGC innervation of the SC

How does visual information from the ipsilateral and contralateral eyes reach binocular neurons in the SC? Contralateral eye input to the SC, carried by RGC axons that cross at the optic chiasm, is well known to terminate in the superficial layers of the mouse SC^41^. In contrast, direct ipsilateral eye input to the SC, originating from uncrossed RGC axons, makes up only 3% of all RGC input to the SC^42^. This weaker ipsilateral eye input to the SC has previously been shown to terminate near the border between the superficial and intermediate layers of the SC^41^. In contrast to the distribution of direct RGC input to the SC, binocular neurons were found in superficial, intermediate, and deep layers of the SC, at depths of up to 1.4 mm from the SC surface (Figure 3B, I). This discrepancy between the location of binocular neurons and previous work on the distribution of direct RGC input in the SC, makes it unclear how binocular SC neurons, particularly those in deeper SC layers, receive visual input from both eyes.

To address this issue, we reinvestigated the precise location of RGC axons in the SC. To do this we studied the anatomical distribution of crossed versus uncrossed RGC axons using publicly available datasets from the Allen Institute for Brain Science. These datasets were comprised of images of fluorescently labeled RGC axons after intravitreal anterograde tracer injections into the contralateral or ipsilateral eye (Figure 4A; see Methods). We found that crossed axons covered almost the entire surface area of the SC (Figure 4B, D), but consistent with earlier work were only found ∼0-450 µm below the dorsal surface of the SC, corresponding to the superficial layers of the SC (Figure 4C, D). In contrast, uncrossed RGC axons were located primarily in a narrow, medial strip that covers the anteroposterior length of the SC, but were strongest in the anterior SC, which is known to processes visual information from the binocular part of the visual field. Uncrossed axons also extended laterally, but only in the most anterior half of the SC. With respect to depth, we found that uncrossed axons were located ∼250-450 µm below the surface of the SC, corresponding to the bottom of the superficial layer. We corroborated these results by performing intravitreal injections with AAV2-Syn-Chronos-GFP (as described below), resulting in comparable crossed and uncrossed RGC axonal expression patterns. Together, these anatomical results indicate that RGC axons innervate only a spatially restricted region of the SC, with overlap of contralateral and ipsilateral eye RGC projections only at the bottom of the superficial layer of the SC.

**Figure 4.**
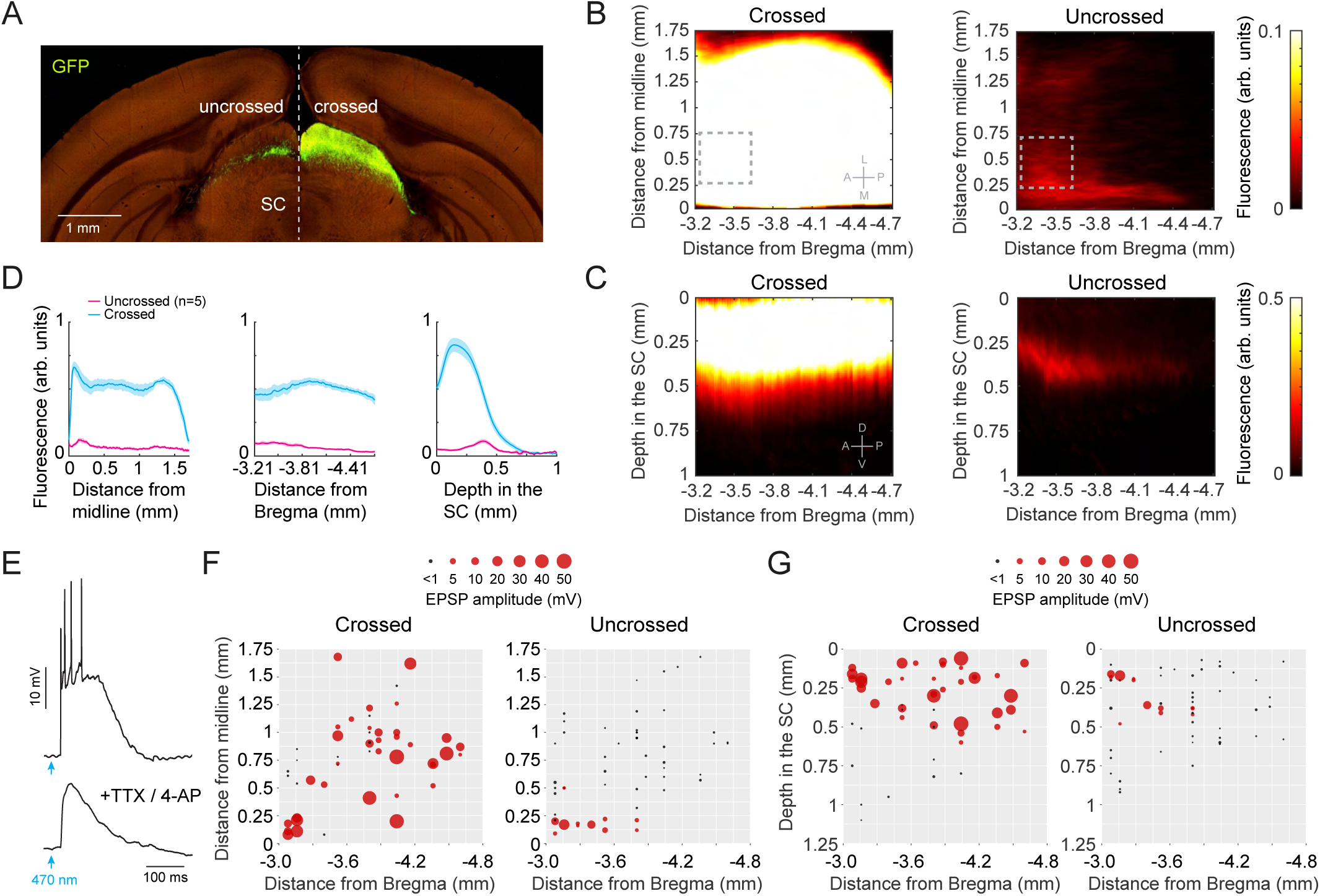
Anatomical and functional characteristics of crossed and uncrossed retinal ganglion cell axons in superior colliculus. (A) Image showing fluorescently-labeled uncrossed (left) and crossed (right) RGC axons in superior colliculus. Taken from the Allen Mouse Brain Connectivity Atlas (connectivity.brain-map.org/projection/experiment/306957248). (B, C) Heat maps showing the average fluorescence of RGC axons along the anteroposterior and mediolateral axes (B) and the anteroposterior and dorsoventral axes (C) of the SC. Fluorescence in heat maps have been capped at the colorbar limit (0.1 or 0.5), with fluorescence exceeding the upper limit shown as white. Gray dotted regions indicate the approximate recording location during *in vivo* recordings. (D) Pooled data summarizing the distribution of RGC axons in the SC from the midline (left), Bregma (middle) and the dorsal surface of the SC (right). Uncrossed RGC axons (pink) localize in the anteromedial SC, mainly between 0.25 and 0.45 mm below the dorsal SC surface. (E) Synaptic response of an SC neuron to optogenetic activation of RGC input (2 ms; 470 nm) in control (top) and TTX plus 4-AP (bottom). (F, G) Distribution of optogenetic response amplitude along the anteroposterior and mediolateral axes (F) and the anteroposterior and dorsoventral axes (G) of the SC. Black responses < 1mV, red responses > 1 mV (circle size indicates response amplitude). Data points show individual neurons. Lines and shaded error bars show mean ± SEM.

### Optogenetic analysis of RGC innervation of the SC

To determine the location of functional synaptic connections between RGCs and SC neurons we performed *in vitro* whole-cell recordings from SC neurons in brain slices from mice expressing AAV2-Syn-Chronos-GFP in only one eye (n=101 neurons from 16 mice). To isolate monosynaptic RGC input, recordings were made from SC neurons in the presence of the sodium channel blocker tetrodotoxin (TTX; 1 µM) and the voltage-gated potassium channel blocker 4-Aminopyridine (Figure 4E; 4-AP; 100 µM). RGC synaptic responses to optogenetic activation (2 ms; 470 nm) in the SC contralateral to the injected eye, receiving crossed RGC input, were observed across the entire SC surface (Figure 4F, left). In contrast, RGC synaptic responses in the SC ipsilateral to the injected eye, receiving uncrossed RGC input, were only found in the most anteromedial region of the SC, consistent with their anatomical localization (Figure 4F; right). In regard to depth, crossed (contralateral eye) RGC synaptic responses were observed in superficial SC layers across the anteroposterior axis, up to ∼0.5 mm from the SC surface (Figure 4G, left). In contrast, uncrossed (ipsilateral eye) RGC synaptic responses were only observed in the most anterior region of the SC at depths between 150 and ∼400 µm from the dorsal SC surface (Figure 4G, right). These data demonstrate that at a functional level convergence of direct RGC input from the contralateral and ipsilateral eye is likely to only innervate SC neurons located at the bottom of the superficial layer of the SC, and only in the most anteromedial part of the SC.

### Tectotectal commissural projections innervate different layers of the opposite SC

These data highlight the discrepancy between the location of direct RGC input to the SC and the location of binocular neurons in the SC, suggesting that binocular visual input to the SC is likely to reach the SC via additional pathways. One possible candidate is from the SC in the opposite hemisphere, as interhemispheric (‘commissural’) projections between the SC in the two hemispheres are known to exist^43–45^. To test this possibility, we unilaterally injected AAV1-hSyn-ChR2(H134R)-EYFP into the SC in one hemisphere (Figure 5A-B), leading to EYFP-labeled axons in the opposite SC (Figure 5C). The density of these commissural projections was highest in the deeper parts of the superficial layer, and the deeper parts of the intermediate and deep layers of the SC (Figure 5D).

**Figure 5.**
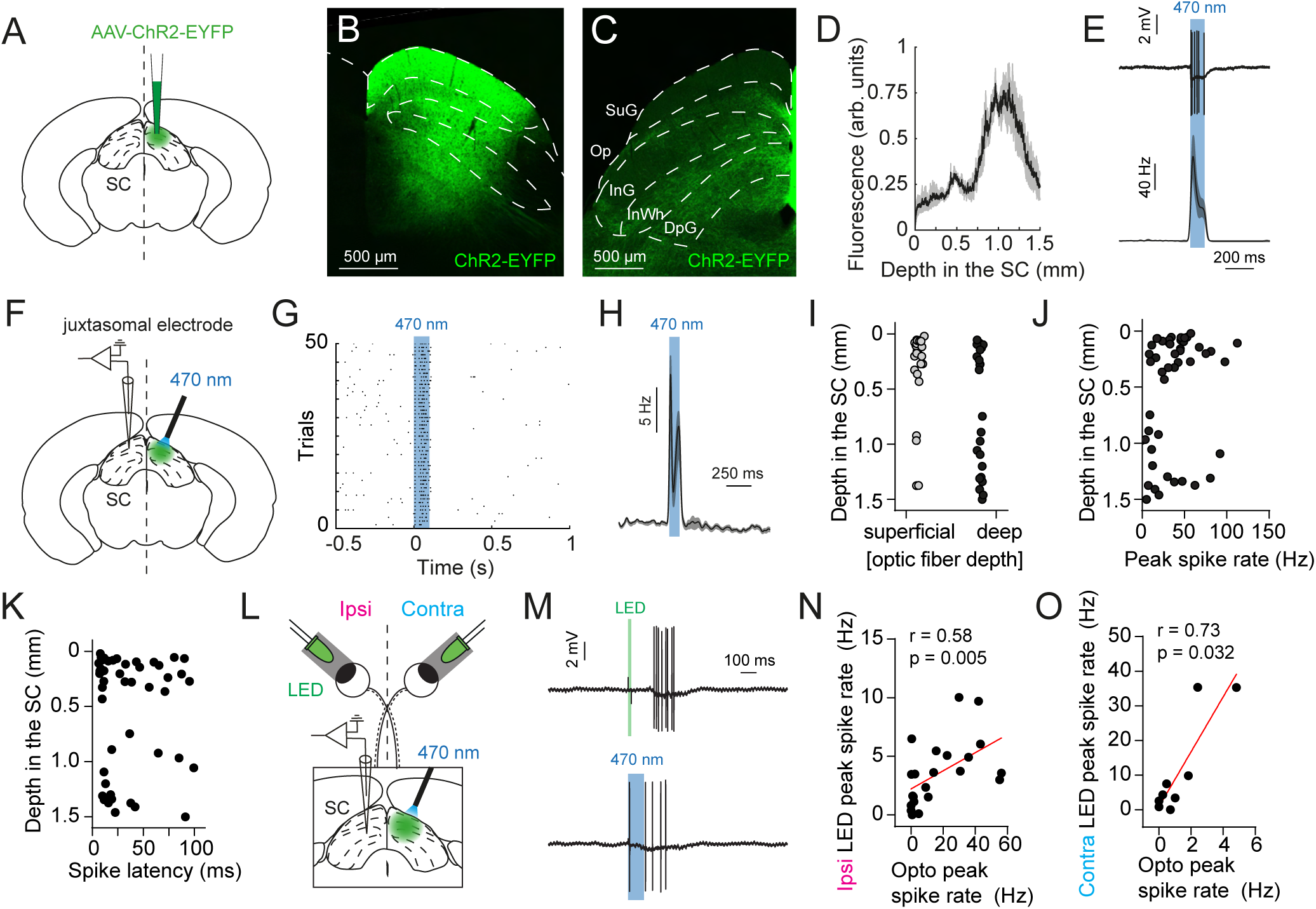
Tectotectal commissural responses correlate with visual responses in superior colliculus. (A) Schematic of unilateral SC viral injection leading to the expression of Channelrhodopsin-2 (ChR2) and enhanced yellow fluorescent protein (EYFP) in one hemisphere of the SC. (B) ChR2-EYFP expression across all layers of the injected SC. (C) SC opposite to the injected SC shown in B. Fluorescently-labeled axons are found in superficial and deeper layers. SuG: superficial gray layer, Op: optic layer, InG: intermediate gray layer, InWh: intermediate white layer, DpG: deep gray layer. (D) Quantification of fluorescence with depth from the SC dorsal surface. (E) Top: Spiking in a ChR2-expressing neuron in the injected SC hemisphere during optogenetic stimulation with 470 nm light (blue bar). Bottom: Average spike rate during optogenetic stimulation (n=3 neurons). (F) Schematic of *in vivo* juxtasomal recordings from the SC opposite to ChR2-EYFP expression. ChR2-expressing neurons were activated via an optic fiber. (G) Peristimulus time histogram of a SC neuron receiving commissural input, showing spike activity during optogenetic stimulation (blue bar). Each dot indicates a spike. (H) Average spiking response (n=24 neurons) to optogenetic stimulation of the opposite SC (blue bar). (I) Depth of neurons receiving commissural input for experiments where the optical fiber for optogenetic activation was superficial (on top of the SC, left) and deep (0.5-0.75 mm below SC surface, right). (J) Peak spike rate as a function of SC depth showing two broad clusters. (K) Spike latency as a function of depth. (L) Schematic of juxtasomal recordings during LED eye stimulation and optogenetic stimulation of the opposite SC via an optical fiber emitting 470 nm light. (M) Spike responses to LED stimulation (top; green bar) and optogenetic stimulation (bottom; blue bar). (N) Peak spike rate during ipsilateral eye and optogenetic stimulation in individual neurons are correlated (r = 0.58, p = 0.005, Spearman, two-tailed). (O) Peak spike rate during contralateral eye and optogenetic stimulation in individual neurons at depths >0.45 mm below SC surface are correlated (r = 0.73, p = 0.032, Spearman, two-tailed). Data points show individual neurons. Lines and shaded error bars show mean ± SEM.

To investigate the functional impact of this pathway, we first confirmed that we could optogenetically activate neurons in the injected SC via an optic fiber positioned above the SC. *In vivo* juxtasomal recordings (n=33 neurons from 3 mice) showed that 470 nm (blue) light could activate ChR2-expressing neurons (Figure 5E; peak spike rate: 109 ± 23.3 Hz, latency: 13.6 ± 4.1 ms). Next, we optogenetically activated these neurons while recording from putative postsynaptic target neurons in the opposite SC (Figure 5F-H; n=109 neurons from 9 mice). Almost half of the recorded SC neurons (44%) increased their spiking during optogenetic activation of the opposite SC (peak spike rate: 37 ± 3.9 Hz, latency: 35.6 ± 4.3 ms). Cells in both superficial and deeper layers of the SC responded to activation of the opposite SC, irrespective of the depth of the optic fiber in the opposite SC (Figure 5I). We found two clusters of SC neurons that responded to commissural input, located ∼0-0.5 mm and ∼0.75-1.5 mm below the dorsal SC surface (Figure 5J-K). These data show that commissural axons make functional connections to SC neurons in superficial and deeper layers of the SC.

### Strength of commissural input correlates with ipsilateral visual response

We next performed *in vivo* juxtasomal recordings (n=22 neurons from 2 mice) during brief LED stimulation of the eyes, while in the same neuron quantifying the strength of commissural input using optogenetic activation of the opposite SC (Figure 5L-M). We found that spiking responses to ipsilateral eye stimulation were correlated with that evoked by optogenetic activation of the opposite SC, indicating that neurons receiving stronger commissural input had stronger ipsilateral visual responses (Figure 5N). A similar correlation between optogenetic and visual-evoked responses was also found for contralateral eye stimulation, but only for neurons located in intermediate and deeper layers (Figure 5O; recording depth >0.45 mm below the SC surface), corresponding to layers of the SC that do not receive direct contralateral RGC input. These results support the idea the commissural pathway from the opposite SC contributes to binocular visual processing in both superficial and deeper layers of the SC.

### Optogenetic silencing of commissural axons decreases binocular responses in SC neurons

To directly test whether the commissural pathway from the opposite SC contributes to binocular visual processing, we optogenetically silenced neurons in the contralateral SC during ipsilateral and contralateral eye stimulation. The SC on one side of Vgat-cre transgenic mice, in which GABAergic neurons express Cre recombinase^46^, was injected with AAV1-EF1a-doublefloxed-hChR2(H134R)-EYFP, leading to cre-dependent ChR2-EYFP expression in SC inhibitory neurons in one hemisphere (Figure 6A-B). *In vivo* juxtasomal recordings from the injected SC (n=19 neurons from 3 mice) indicated that optogenetic stimulation led to spiking in putative Vgat+ neurons, and inhibition of spiking in putative Vgat-neurons (Figure 6C). We next made *in vivo* juxtasomal recordings from Vgat-neurons during contralateral eye stimulation with or without optogenetic activation of Vgat+ neurons (Figure 6D-E). This led to a 90 ± 6% decrease in spiking evoked by brief LED stimulation of the contralateral eye (Figure 6F). These data confirm that optogenetic activation of inhibitory neurons in one hemisphere of the SC is an effective method to silence visual responses in that hemisphere, and thereby block the transfer of visual input to the opposite SC via the commissural pathway.

**Figure 6.**
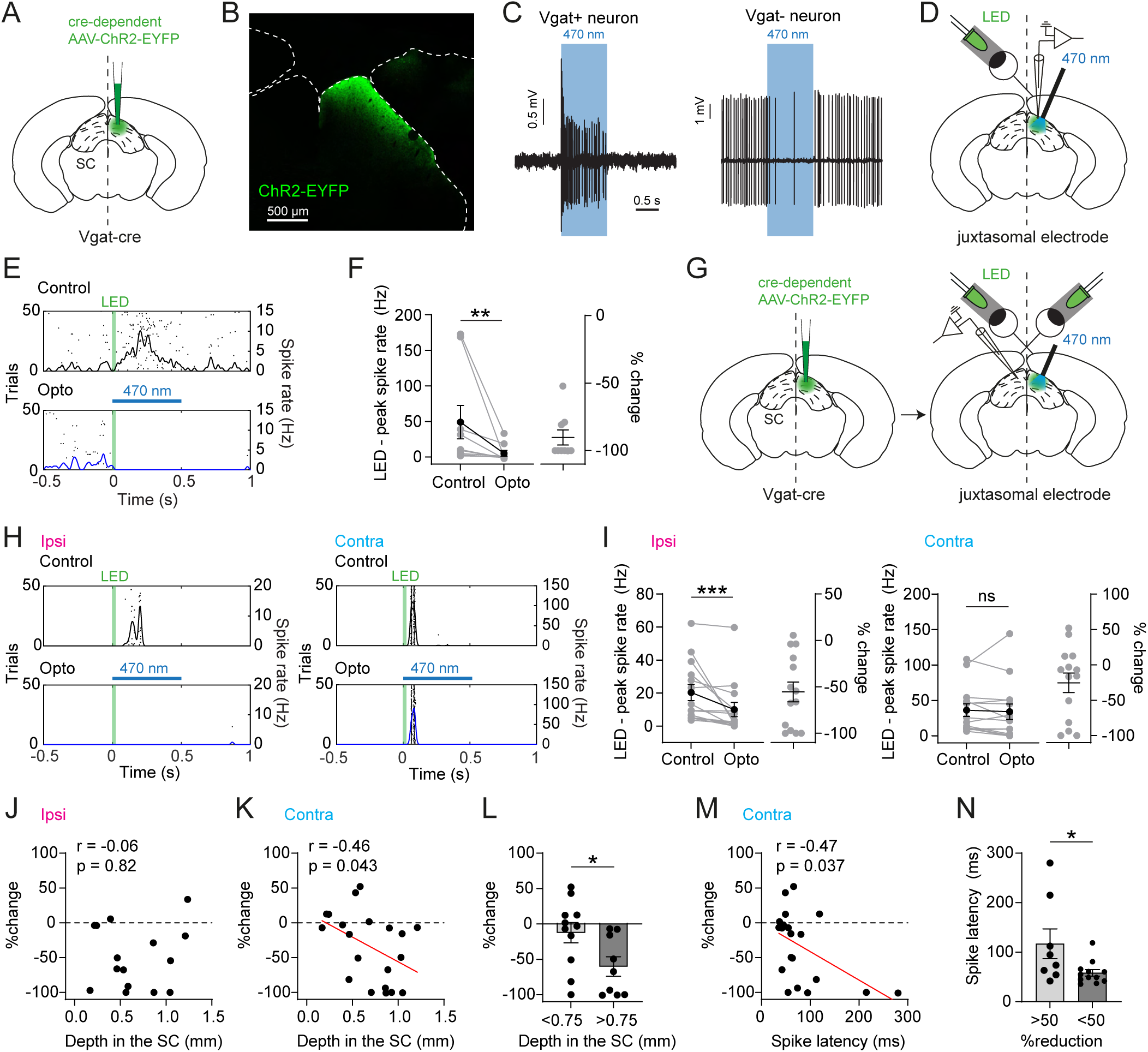
Silencing commissural projections reduces ipsilateral and contralateral visual responses in superior colliculus. (A) Schematic showing unilateral cre-dependent ChR2 viral injection into one hemisphere of the SC in Vgat-cre mice. (B) Viral injection led to cre-dependent expression of Channelrhodopsin-2 (ChR2) and enhanced yellow fluorescent protein (EYFP) in SC neurons only in one hemisphere. (C) Spiking responses from putative Vgat+ and Vgat- neurons, showing an increase and a decrease, respectively, in spiking during optogenetic stimulation (blue bar). (D) Schematic of *in vivo* juxtasomal recording from neurons in the injected SC during contralateral eye stimulation. ChR2-expressing neurons were optogenetically activated by an optical fiber emitting 470 nm light. (E) Peristimulus time histograms showing spiking during contralateral eye stimulation without (‘control’, top) and with optogenetic silencing (‘opto’, bottom). Each dot shows a spike and lines show spike rate (right Y-axis). (F) Impact of optogenetic silencing on contralateral eye spiking in individual neurons. Right: Percentage change relative to the control response (n=9 neurons from 3 mice; control vs. opto: W = -45, p = 0.004, Wilcoxon matched-pairs signed rank test, two-tailed). (G) Schematic showing unilateral virus injection in the SC of Vgat-cre mice (right) together with *in vivo* juxtasomal recording in the opposite SC during ipsilateral and contralateral eye stimulation with LEDs and optogenetic stimulation by an optical fiber emitting 470 nm light. (H) Peristimulus time histograms showing spiking during ipsilateral (left) and contralateral (right) eye stimulation (green bars) without (‘control’, top) and with optogenetic silencing of the opposite SC (‘opto’, bottom, blue bars). Each dot shows a spike and lines show estimates of spike rate (right Y-axis). (I) Impact of optogenetic silencing of the opposite SC on ipsilateral (left) and contralateral (right) eye spiking in individual binocular SC neurons. Percentage change relative to the control response shown on the right. Optogenetic stimulation decreases ipsilateral, but not contralateral, eye-evoked spike responses (ipsi: W = -101, p < 0.001, Wilcoxon matched-pairs signed rank test, two-tailed; contra: W = -45, p = 0.173, Wilcoxon matched-pairs signed rank test, two-tailed). (J-K) Percentage change in ipsilateral (J; r = -0.06, p = 0.82, Spearman, two-tailed) and contralateral (K; r = -0.46, p = 0.043, Spearman, two-tailed) eye-evoked spiking in SC neurons during optogenetic silencing versus their depth in the SC. (L) Percentage change in contralateral eye-evoked responses during optogenetic silencing for superficial (<0.75 mm depth) versus deep (>0.75 mm depth) SC neurons (*t*(17.91) = 2.394, p = 0.028, Welch’s t-test). (M) Percentage change in contralateral eye-evoked spiking in SC neurons during optogenetic silencing versus the eye-evoked spike latency (r = -0.47, p = 0.037, Spearman, two-tailed). (N) Eye-evoked spike latency separating neurons showing >50% versus a <50% change in contralateral eye-evoked responses during optogenetic stimulation (U = 22, p = 0.047, Mann-Whitney U-test). Data points show individual neurons. Lines with error bars show mean ± SEM. *p < 0.05, **p < 0.01, ***p < 0.001.

Next, we performed *in vivo* juxtasomal recordings from binocular neurons in the SC opposite to that expressing cre-dependent ChR2-EYFP during brief LED eye stimulation of the contralateral and ipsilateral eyes with or without optogenetic silencing of the opposite SC (Figure 6G-H; n=14 neurons from 6 mice). We only included neurons in this analysis that did not change their spontaneous spike rate during optogenetic stimulation alone. Across all binocular SC neurons, we found that silencing the opposite SC led to a 56 ± 11% decrease in the ipsilateral eye-evoked response, but no change in contralateral eye-evoked responses (Figure 6I). However, the percentage change in contralateral, but not ipsilateral, eye-evoked responses during optogenetic silencing of the opposite SC was correlated with cell depth in the SC (Figure 6J-K). Further analysis indicated that contralateral eye responses were significantly reduced by optogenetic silencing of the opposite SC for neurons in intermediate and deeper layers (>0.75 mm depth), whereas this was not the case for neurons in more surficial layers of the SC (Figure 6L). We also found a correlation between the percentage change in the contralateral eye-evoked response following optogenetic silencing of the opposite SC and spike latency (Figure 6M). Furthermore, SC neurons showing more than 50% reduction in the contralateral eye responses had significantly longer spike latencies, suggesting that polysynaptic pathways may carry contralateral eye input to these neurons (Figure 6N). Together, these experiments demonstrate that interhemispheric commissural input conveys both ipsilateral and contralateral eye information to the opposite SC. While commissural ipsilateral eye input is observed across all SC layers, contralateral eye-evoked responses mediated via interhemispheric commissural input are only observed in intermediate and deeper SC layers.

### Contralateral V1 conveys binocular visual input to the SC

The SC also receives prominent input from layer 5 pyramidal neurons in V1^18,23^, which have been previously shown to modulate SC visual responses *in vivo*^47^. While most V1 neurons are driven by contralateral visual input, a subset of neurons in V1 respond to both ipsilateral and contralateral eye input and are therefore binocular^32,33,48^. We hypothesized that these binocular layer 5 neurons may convey binocular visual input to SC neurons. In rodents, binocular neurons in V1 are located in a narrow strip at the border between areas 17 and 18, which receives callosal input from the opposite V1^49,50^. To investigate whether SC-projecting V1 neurons are located in this area, we labeled V1 callosal axons with AAV1-hSyn-ChR2(H134R)-EYFP and retrogradely labeled SC-projecting V1 neurons by injecting AAV2-retro-EF1a-mCherry-Cre into the SC (Figure 7A). SC-projecting layer 5 pyramidal neurons in V1 (mCherry+) were found within the terminal region of EYFP-expressing V1 callosal projections (Figure 7B), consistent with the idea that SC projecting V1 neurons could be binocular, and therefore may convey both contralateral and ipsilateral eye input to the SC.

**Figure 7.**
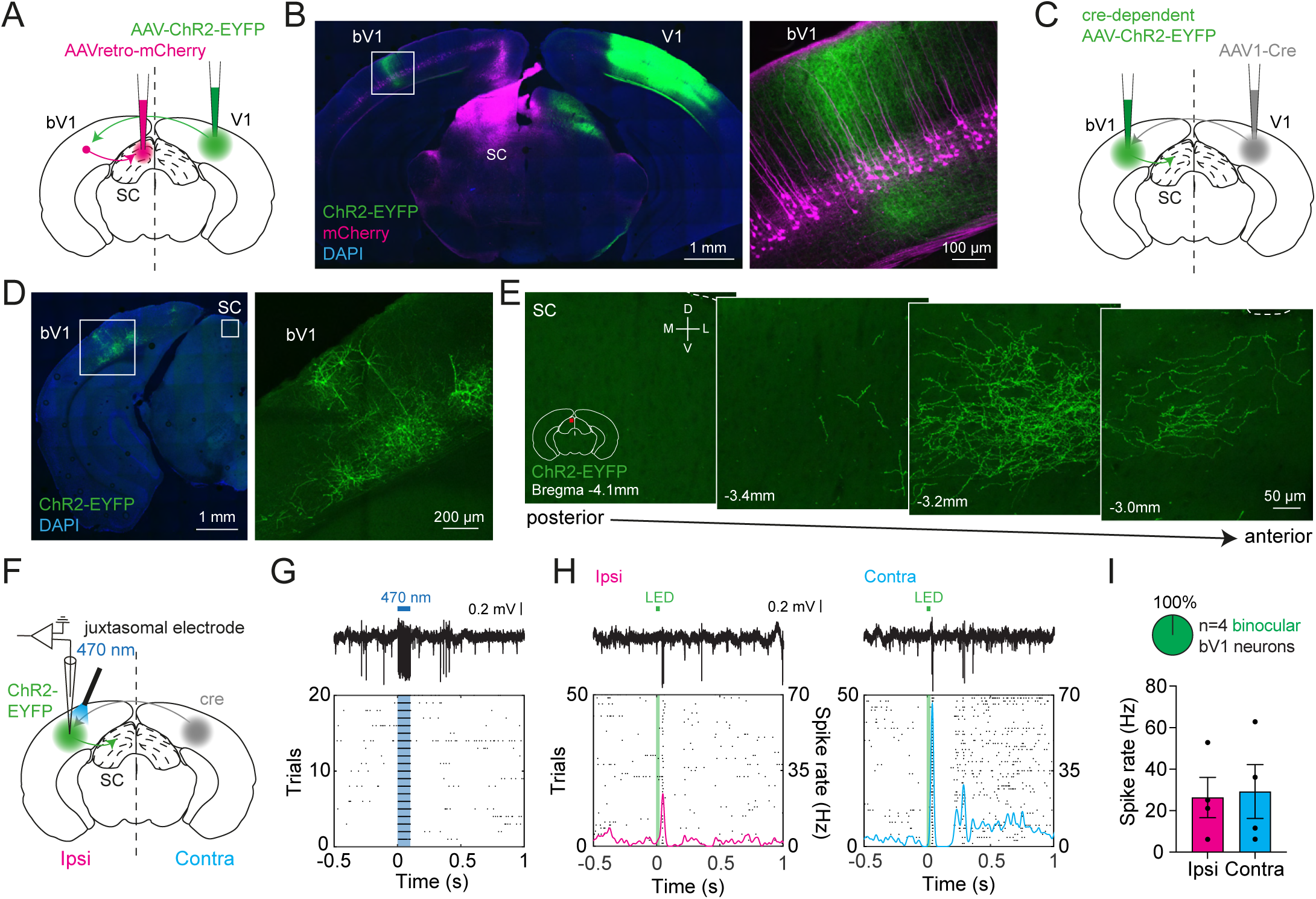
Binocular primary visual cortex projects to anteromedial superior colliculus. (A) Schematic of AAV-ChR2-EYFP injection into the right V1 and AAVretro-mCherry injection in the left SC. bV1: binocular area in V1. (B) Expression of EYFP in commissural axons demarcating binocular V1. mCherry-expressing cell bodies of SC-projecting layer 5 pyramidal neurons are located in binocular V1. White square in the left image indicates the position of the magnified inset on the right. (C) Schematic of AAV1-Cre injection into the right V1 and cre-dependent AAV-ChR2-EYFP into the left SC. Trans-synaptic transfer of cre enables postsynaptic expression of ChR2-EYFP in commissural receiving V1 neurons. (D) ChR2-EYFP expression in commissural target neurons in the left binocular V1. Neurons are present in multiple layers, including layer 5. White square in the left image indicates the position of the magnified inset on the right. (E) Expression of ChR2-EYFP in axons from neurons receiving commissural input in the SC. Expression is centered around -3.2 mm Bregma. Imaging location is indicated by the red square in the schematic drawing. D: dorsal, V: ventral, M: medial, L: lateral. (F) Schematic of juxtasomal recording from the left binocular V1, where neurons receiving commissural input express ChR2-EYFP. Optogenetic activation of these neurons was achieved by an optic fiber emitting 470 nm light. (G) Top: Spiking of a ChR2-EYFP expressing V1 neuron during optogenetic stimulation (blue bar). Bottom: Peristimulus time histogram showing spike activity on different trials. Each dot shows a spike. (H) Top: Spiking in the same neuron shown in G during ipsilateral (left) and contralateral (right) eye stimulation using brief LED illumination (green). Bottom: Peristimulus time histograms showing spike activity on different trials. Each dot shows a spike. Colored lines show spike rate (right Y-axis). (I) Spike rate during ipsi- and contralateral eye stimulation in optogenetically-identified V1 neurons receiving commissural input, showing that these neurons are binocular. Data points in I show individual neurons. Bar plots show mean ± SEM.

Next, to investigate whether binocular V1 neurons project to the SC, we took advantage of AAV1-mediated trans-synaptic labeling methods^28^. As only binocular V1 neurons receive callosal projections from the opposite V1^51^, we injected AAV1-hSyn-Cre into V1 in one hemisphere, followed by AAV1-EF1a-doublefloxed-hChR2(H134R)-EYFP into V1 in the opposite hemisphere, leading to cre-dependent ChR2-EYFP expression in binocular V1 neurons receiving callosal input (Figure 7C-D). We then investigated where these binocular V1 neurons project, finding EYFP-labeled axons in the anteromedial region of the SC, at a location where visual information from the upper binocular visual field is processed (Figure 7E). To verify that labelled V1 neurons are binocular we performed juxtasomal recordings from ChR2-expressing V1 neurons, identified by short-latency responses to optogenetic light activation (Figure 7F-G). All recorded neurons showed spiking responses to brief ipsi- and contralateral eye LED stimulation, indicating they are indeed binocular (Figure 7H-I). Together, these data support the idea that binocular visual input to the SC can arise from SC-projecting binocular V1 neurons.

### Optogenetic silencing of binocular V1 neurons leads to decreased ipsilateral eye responses in the SC

To determine whether the binocular V1 corticotectal pathway conveys visual input to the SC, we injected AAV5-Syn-FLEX-rc[ChrimsonR-tdTomato] into binocular V1 of Vgat-cre mice, leading to cre-dependent expression of the excitatory opsin ChrimsonR in inhibitory GABAergic neurons in binocular V1 (Figure 8A-B). This enabled us to optogenetically activate inhibitory neurons in binocular V1 using 590 nm light (Figure 8C). Recordings from putative vGat-layer 5 pyramidal neurons in the injected binocular V1 (n=12 neurons from 2 mice) confirmed that visually-evoked spike responses were reduced by on average 73 ± 10% during ChrimsonR activation (Figure 8D). Next, we performed juxtasomal recordings from binocular neurons in the SC in the same hemisphere (n=8 neurons from 4 mice) during brief LED stimulation of the ipsi- or contralateral eyes, with or without optogenetic silencing of binocular V1 on alternating trials (Figure 8E). Silencing of binocular V1 reduced ipsilateral eye spiking responses by on average 64 ± 15%. Neurons showing the largest reduction in firing were located around 0.5 mm from the SC surface (Figure 8F). Somewhat surprisingly, we did not observe a significant reduction in contralateral eye responses (Figure 8G). Consistent with this spiking data, using *in vivo* whole-cell recording we found that V1 silencing reduced synaptic responses evoked by stimulation of the ipsilateral, but not the contralateral, eye (Figure S5). These data confirm that the corticotectal pathway from binocular V1 conveys visual information to binocular neurons in the SC, contributing primarily to the ipsilateral eye response.

**Figure 8.**
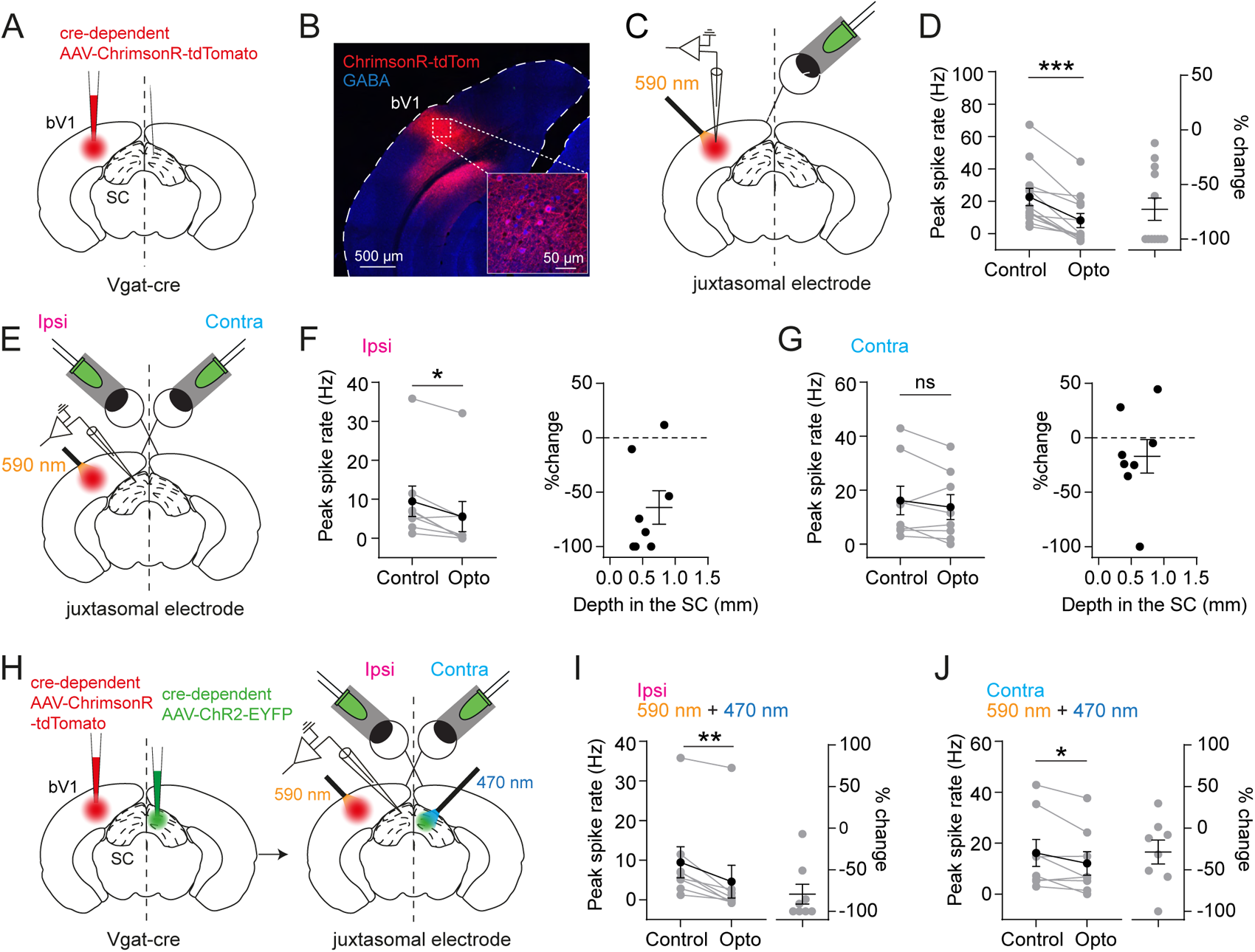
Effect of silencing visual pathways through V1 on responses in superior colliculus. (A) Schematic of unilateral virus injection into Vgat-cre mice leading to cre-dependent expression of ChrimsonR-tdTomato in binocular V1 (bV1). (B) Expression of ChrimsonR-tdTomato in GABAergic neurons across all layers of the injected binocular V1. (C) Schematic of *in vivo* juxtasomal recordings from the injected binocular V1. ChrimsonR-tdTomato-expressing neurons are activated by an optic fiber emitting 590 nm light and stimulation of the contralateral eye using a LED. (D) Contralateral eye spiking was reduced by optogenetic activation of Vgat neurons (W = -78, p < 0.001, Wilcoxon matched-pairs signed rank test, two-tailed). (E) Schematic of *in vivo* juxtasomal recordings from the SC ipsilateral to the injected binocular V1. ChrimsonR-tdTomato-expressing neurons are activated by an optic fiber emitting 590 nm light and the ipsilateral and contralateral eyes via LEDs. (F-G) Ipsilateral eye (F) but not contralateral eye spiking responses (G) in binocular SC neurons are reduced by optogenetic silencing of binocular V1 (ipsi: W = -34, p = 0.016, Wilcoxon matched-pairs signed rank test, two-tailed; contra: W = -20, p = 0.195, Wilcoxon matched-pairs signed rank test, two-tailed). (H) Schematic showing injections of cre-dependent AAV-ChrimsonR-tdTomato in left binocular V1 and cre-dependent AAV-ChR2-EYFP in the opposite SC in Vgat-cre mice. Injections are followed by *in vivo* juxtasomal recordings from the SC. ChrimsonR-tdToma-to-expressing neurons are activated using 590 nm light, ChR2-EYFP-expressing neurons by 470 nm light and visual stimulation of the ipsilateral and contralateral eyes using LEDs. (I-J) Dual silencing leads to a strong decrease in ipsilateral eye-evoked responses (I; W = -36, p = 0.004, Wilcoxon matched-pairs signed rank test, one-tailed) and contralateral eye-evoked responses (J; W = -26, p = 0.039, Wilcoxon matched-pairs signed rank test, one-tailed). Data points show individual neurons. Lines and error bars show mean ± SEM. *p < 0.05, **p < 0.01, ***p < 0.001.

### Dual optogenetic silencing of binocular V1 and the opposite SC significantly impacts binocular processing in SC neurons

In a final set of experiments, we determined the impact of silencing of both the ipsilateral binocular V1 and the opposite SC on binocular responses in the SC. To investigate this, Vgat-cre mice were co-injected with AAV5-Syn-FLEX-rc[ChrimsonR-tdTomato] in binocular V1 and AAV1-EF1a-doublefloxed-hChR2(H134R)-EYFP in the opposite SC, resulting in the expression of ChrimsonR and ChR2 in GABAergic neurons in binocular V1 and the opposite SC, respectively (Figure 8H, left). To activate each opsin independently we placed a 590 nm light-emitting optic fiber above binocular V1 and a 470 nm light-emitting optic fiber above the opposite SC. *In vivo* juxtasomal recordings were then made from SC neurons ipsilateral to ChrimsonR expression in binocular V1 and contralateral to ChR2 expression in the opposite SC (Figure 8H, right). Binocular responses were evoked by ipsilateral and contralateral eye stimulation using brief LED flashes with or without optogenetic silencing of binocular V1 and the opposite SC on alternating trials. This revealed that ipsilateral eye responses were severely decreased by 79 ± 12% during combined silencing of binocular V1 and the opposite SC (Figure 8I; n=8 neurons from 4 mice). Contralateral eye responses were also decreased, but to a smaller extent (29 ± 14%; Figure 8J). Together these findings highlight that both the tectotectal and corticotectal commissural pathways play an important role in conveying binocular visual input to the SC.

## Discussion

In this study, we shed light on the importance of binocular vision for defensive behaviors to visual threats. Furthermore, we describe the synaptic and spike mechanisms underlying binocular integration in the SC, and provide a detailed analysis of the pathways that convey binocular visual information to SC neurons. Importantly, we show that disrupting binocular vision through monocular enucleation led to impaired escape responses and increased freezing, without affecting their ability to detect threatening visual stimuli. Since the SC is thought to play a crucial role in orchestrating escape and freezing responses to threatening visual stimuli^5^, we focused our investigation on binocular processing in the SC. *In vivo* synaptic and spike recordings revealed a variety of synaptic and spike response patterns to binocular stimuli, consistent with recent studies^34,35^. To determine which neural pathways are responsible for conveying visual input to SC neurons, we characterized RGC axonal projections to the SC from the contralateral and ipsilateral eye and found that they innervate spatially restricted regions in the superficial SC layer, with ipsilateral eye input restricted to the bottom of the superficial layer in the anteromedial part of the SC. Consistent with this anatomical analysis, functional synaptic connections were found in the same SC regions. Given that binocular neurons in the SC were found across superficial, intermediate and deep layers, these observations suggested that binocular visual input to the SC is likely to be mediated via multiple visual pathways. Consistent with this idea, using both anatomical labelling and optogenetic silencing we show that in addition to direct input from the retina, binocular visual input is also conveyed to the SC via interhemispheric (‘commissural’) projections from the opposite SC as well as ipsilateral binocular V1.

### Limitations

There are several factors that may have impacted our results. Firstly, regarding the behavioral experiments it should be noted that monocular enucleation does not only lead to loss of binocular information, but also to a loss of perception in one half of the visual field. This did not significantly impact the capacity of mice to detect threatening visual stimuli, however, since we did not observe an increased number of miss trials in enucleated mice. Secondly, despite significant reductions in visual responses in our optogenetic silencing experiments, optogenetic silencing of the opposite SC and/or binocular V1 did not completely block visual responses. This may simply result from incomplete silencing of the tectotectal or corticotectal pathways. If so, the contribution of these pathways to binocular visual responses in the SC may have been underestimated. Conversely, as commissural tectotectal projections are both excitatory and inhibitory^45,52^, activation of Vgat+ neurons during silencing the opposite SC may inadvertently excite inhibitory neurons projecting to the opposite SC. This could lead to increased inhibition in the opposite SC, reducing eye-evoked responses. If this occurred, we may have overestimated the impact of tectotectal projections on eye-evoked responses. We do not think this is the case because we only included neurons in our analysis that did not show a change in spontaneous spiking during optogenetic silencing of the opposite SC. Finally, we did not find an impact of silencing ipsilateral V1 on contralateral eye responses. This is somewhat surprising given the strong projection of V1 to SC, and our finding that silencing V1 led to a reduction in ipsilateral eye responses. One possible explanation for this is that contralateral eye responses in the SC, at least in the superficial layers, are dominated by direct, crossed RGC input, which as we show is much more powerful than uncrossed RGC input from the ipsilateral eye. In this regard, it is worth noting that the recordings in these experiments were in the upper half of the SC. Another possibility is that a reduction in V1 excitatory drive to inhibitory neurons in the SC may have led to a local increase in excitation (disinhibition), masking the impact of V1 silencing on contralateral eye responses.

### Modulation of freeze and escape behavior selection

Recent studies have uncovered that the SC mediates innate defensive behaviors through its projections to downstream regions such as the LP, PBG and PAG^3,4,25,28,53^. Optogenetic activation of SC-PBG projections results in escape, while activation of SC-LP projections results in freezing^3^, suggesting that threatening visual input may initiate distinct defensive responses depending on the spatiotemporal characteristics of the visual threat^1,3,5^. While this simplistic description of defensive behaviors is appealing, the selection of defensive behaviors is also influenced by a range of factors including stress^54^, position relative to a shelter^55^, preceding events^56^ and top-down or bottom-up influences from other brain regions^57–59^. The “decision” to escape or freeze in response to a threatening visual stimulus is therefore influenced by multiple factors. Our results indicate that when binocular vision is disrupted, there is a shift towards freezing rather than escape, indicating that binocular vision also contributes to the decision whether to escape or freeze in response to visual threat. A shift towards increased freezing is also observed during other manipulations of SC activity, such as optogenetic activation of input from V1^28^.

One interpretation of our observations is that reduced activity in binocular SC neurons leads to an imbalance between PBG and LP activation, resulting in an increase in freezing behavior. Consistent with this idea, three of the four main cell types in the superficial SC project to the PBG, *i.e.,* stellate, horizontal and narrow-field cells, whereas only one cell type projects to LP, *i.e.* wide-field (WF) cells^60^. As our results suggest that all four SC cell types in the superficial SC are binocular, reducing activation of binocular SC neurons, as is likely to occur in enucleated mice, may lead to a greater reduction in activation of PBG compared to LP. This may tip the scales, so to speak, in favor of freezing rather than escape.

### Visual inputs shape binocular activity in SC

Binocular processing in the mouse SC has only recently been studied^34,35^, with earlier work on binocular processing in the SC investigated primarily in cats and hamsters^61–65^. A commonality between this recent work in mice and our study is that binocular responses in the SC are diverse and more complex than binocular responses in mouse V1^33^. Ipsilateral and contralateral eye stimulation could induce both excitatory and inhibitory synaptic responses, even within the same SC neuron. This could be due to local network interactions. In the superficial SC horizontal neurons are thought to be inhibitory and connect to adjacent cell types^37,60,66^, but inhibitory inputs may also come from commissural tectotectal projections^45,52^. Excitatory connections between superficial and deeper layers of the SC also exist^60^. These intricate network interactions are likely to contribute to the complexity of binocular responses observed, reflecting a dynamic interplay of excitatory and inhibitory mechanisms shaping binocular processing in the SC. Another common finding between our study and recent studies^34,35^ is that contralateral visual input dominates the binocular response. Despite this, ipsilateral visual input influenced binocular responses, leading to sublinear integration during binocular stimulation. This was the case during LED-flash stimuli as well as during spatiotemporally more complex looming stimuli. Both types of stimuli are behaviorally relevant, as LED flashes induce SC-dependent arrest behavior^27^ and looming spots evoke SC-dependent defensive behaviors^2,3^. One possible explanation for why sublinear integration was observed in binocular SC neurons during stimulation of both eyes together is that this was due to recruitment of inhibition, as is the case in V1^33,38^.

### Multiple complementary visual pathways contribute to binocular activity in SC

In terms of visual pathways, RGC inputs have long been thought to be the primary pathway by which visual input is conveyed to SC, particularly for ipsilateral eye input^6,42^. Since both crossed and uncrossed RGC inputs innervate spatially and layer-restricted regions of the SC, we hypothesized that alternative visual pathways are responsible for the functional innervation of binocular SC neurons in regions of the SC that do not receive direct RGC input. Our experiments revealed that two complementary pathways, the tectotectal and corticotectal pathways relay ipsilateral and contralateral visual information to the SC in a layer-specific manner. Silencing the opposite SC had a severe impact on ipsilateral eye responses, and on contralateral visual responses of neurons in deeper layers of the SC. Silencing ipsilateral binocular V1 also led to a strong decrease in ipsilateral eye responses, although had little impact on contralateral eye response. Together, our findings indicate that contralateral and ipsilateral visual information reaches binocular neurons via multiple pathways, which target different layers of the SC. Given that the different layers of the SC are likely to process different streams of sensory information, our data suggests that these different binocular visual pathways to the SC have different functional roles.

In conclusion, our study highlights the importance of binocular visual processing in the SC for innate defensive behaviors, showing that binocular input to the SC is mediated by multiple visual pathways in a layer specific manner. Further work will need to be done to determine how these different binocular visual pathways work together to orchestrate an appropriate defensive response to visual threat.

## Acknowledgments

This work was supported by the National Health and Medical Research Council of Australia, and the Australian Research Council Centre of Excellence for Integrative Brain Function (ARC Centre Grant CE140100007). We thank W.M. Connelly, H. Huang, S. Honnuraiah and S. Gharaei for their contribution to the study. We thank N. Dehorter and N. Ahmed for kindly providing the PV-cre::tdTomato mice and C. Dayas, L. Manning and L. Greco for kindly providing the Vgat-ires-cre mice. We thank S. Voerman and C.I. De Zeeuw for their comments on the manuscript. Graphical abstract created with BioRender.com.

## Author contributions

Conceptualization, R.B. and G.J.S.; Methodology, R.B. and G.J.S.; Investigation, R.B., G.T. and F.T.; Analysis, R.B., G.T. and F.T.; Writing – Original Draft, R.B. and G.J.S.; Writing – Review & Editing, R.B., G.T., F.T. and G.J.S; Funding Acquisition, G.J.S.; Supervision, R.B. and G.J.S.

## Resource availability

### Lead contact

Further information and requests for resources and reagents should be directed to and will be fulfilled by the lead contact, Greg J. Stuart

### Materials availability

This study did not generate new unique reagents.

### Data and code availability

All code used in this article will be available at the time of manuscript acceptance via GitHub https://github.com/BroRobin/BinocSC/

### Data availability statement

All data generated during this study is unreservedly available upon request from the corresponding authors.

## Methods

### Animals and ethics statement

Adult male and female C57BL/6 mice (4-12 weeks of age) were used in this study. Mice were socially housed were possible, with *ad libitum* access to food and water. They were kept in a controlled environment with a 12/12 h light/dark cycle. Experiments were carried out during the light phase. All animal experimental procedures were approved by the Animal Experimentation Ethics Committee of the Australian National University and carried out in accordance with the National Health and Medical Research Council (NHMRC) Australian code for the care and use of animals for scientific purposes.

### Transgenic mouse lines

Vgat-ires-cre mice [B6J.129S6(FVB)-Slc32a1^tm2(cre)Lowl^/MwarJ, Jackson Laboratories, ME, USA; stock number #028862]^46^, expressing Cre in vGAT-expressing neurons, were obtained from Dr. Christopher V. Dayas (University of Newcastle). PV-cre mice [B6;129P2-Pvalb^tm1(cre)Arbr^/J, Jackson Laboratories, ME, USA; stock number #008069]^67^, expressing Cre in Parvalbumin-expressing neurons, were obtained from Dr. Nathalie Dehorter (Australian National University). In some experiments heterozygous PV-Cre::Td-Tomato mice were used, which were generated by crossing PV-cre mice and Ai9 mice [B6.Cg-Gt(ROSA)26Sor^tm9(CAG-tdTomato)Hze^/J, Jackson Laboratories, ME, USA; stock number #007909]^68^.

### Viral vectors

Viruses were obtained from the viral vector cores of the University of Pennsylvania (Penn Vector Core, PA, USA), University of North Carolina (UNC Vectore Core, NC, USA) and Addgene (MA, USA). Viruses were injected in wild type or transgenic mice as indicated in the Results. The viral vectors used were: AAV1-hSyn-ChR2(H134R)-EYFP.WPRE.hGH (Penn Vector Core), AAV1-EF1a-doublefloxed-hChR2(H134R)-EYFP-WPRE-HGHpA (#20298-AAV1, Addgene), AAV2-retro-EF1a-mCherry-IRES-Cre (#55632-AAVrg, Addgene), AAV1-hSyn-Cre-WPRE.hGH (Penn Vector Core), AAV5-Syn-FLEX-rc[ChrimsonR-tdTomato] (#62723-AAV5, Addgene), AAV2-Syn-Chronos-GFP (UNC Vector Core). All viruses were stored at -80 °C prior to use.

### Enucleation

Mice (male/female, 3-5 weeks of age) were anesthetized with isoflurane (3.5% induction; 1.8-2.0% maintenance in 0.3 L/min O_2_), placed on a temperature-controlled heating pad (37 °C; Harvard Instruments) and mounted in a stereotaxic apparatus (Kopf). Depth of anesthesia was monitored frequently by testing for the loss of the withdrawal reflex to a paw pinch. Both eyes were moisturized using ophthalmic gel (GenTeal gel, Alcon) and the analgesic Meloxicam (Metacam, 5 mg/kg s.c.) was injected subcutaneously for pain relief. The left eyeball was displaced by gently pressing onto the canthus using forceps (curved #3, Dumond) and a knotted thread was placed around the eye. The knot was tightened around the posterior bundle containing the optic nerve and arteries, after which forceps were used to remove the eye. A drop of antiseptic solution (Betadine) was topically applied to the eye socket. Mice were kept warm on a heating pad until fully recovered and monitored closely post-surgery. Mice were allowed to recover for at least two weeks before being used for behavioral experiments.

### Open field arena

An open field arena (48cm(l) x 38cm(w) x 30 cm(h)), constructed from white acrylic plastic^1^, was used to measure behavioral responses to threatening visual stimuli. An opaque triangular refuge (7cm(h) x 8cm(w)) was placed in one corner of the arena to provide shelter. A LCD display monitor (60 Hz, 38cm (l) x 27 cm (w)), connected to a computer, was placed on top of the arena and used to present visual stimuli. A square ‘start’ zone (8cm x 8cm) located under the center of the monitor was indicated on the floor of the arena with four black dots for each corner. A high-speed camera (PlayStation 3 Eye) was used to record mouse movements. It was connected to a computer running acquisition software (OBS Studio) and placed so that it could observe the arena floor through the space between the monitor and the adjacent wall. Videos were captured at 75 Hz and saved in QuickTime (.mov) format for offline analysis. A red LED light which was active during presentation of visual stimuli was placed on the upper wall of the arena to aid synchronization of behavioral analysis. It could be detected by the camera but not seen by the mouse.

### Measurements of innate defensive behaviors

Behavioral responses to threatening visual stimuli were recorded over the course of six days. Mice were habituated on day 1 by placing them in the experimental arena for 15 minutes. Background light from a gray screen on the LCD monitor illuminated the arena, but no stimuli were presented. On day 2-6 mice were first allowed to explore the arena for at least 5 minutes, after which two visual stimulus protocols were run with an inter-trial interval (ITI) of at least 3.5 minutes. The visual stimuli used were a sweeping stimulus and a looming stimulus. These stimuli were presented on a light gray background. The sweeping stimulus consisted of a 2.5 cm (5 visual degrees) black circle moving diagonally from one corner to the opposite corner of the monitor over 4 s (∼11 cm/s speed). The looming stimulus was a 1 cm (2 visual degrees) black circle that rapidly widened to 25.5 cm (50 visual degrees) over 250 ms, after which it remained stationary for another 500 ms. These stimuli have previously been shown to induce strong innate defensive behaviors^1^. Stimuli were generated in MATLAB (R2020a, MathWorks) using Psychtoolbox-3^69^ on a dedicated high-performance Windows-based computer and were manually started when the mouse entered the central ‘start’ zone.

### Analysis of innate defensive behaviors

Mouse position was tracked semi-automatically using a modified version of the ‘Mouse Activity Analyzer’ written by Dr. Renzhi Han (The Ohio State University). Coordinates were obtained by pixel-thresholding video images and fitting a centroid area to pixel clusters representing the mouse, to detect objects using the ‘regionprops’ function in MATLAB. Mouse speed was calculated from centroid coordinates on a frame-by-frame basis, converted to distance and filtered with a moving median filter (53.3 ms window; 4 frames). Classification of responses, and registering their timing relative to stimulus onset, was done manually with the observer blind to the experimental group analyzed. Responses were classified into five categories: escape, freeze, escape and freeze, no response or under refuge. Escape was defined as a sudden and rapid movement towards the refuge or periphery of the arena (Video S1). Freezing was defined as a complete cease of motion for at least 0.5 s.

### Intracranial viral injections

Mice (male/female, 3-5 weeks of age) were anesthetized with isoflurane (3.5% induction; 1.8-2.0% maintenance in 0.3 L/min O_2_), placed on a feedback-controlled heating pad set to 37 °C (Harvard Instruments) and mounted in a stereotaxic apparatus (Kopf). Depth of anesthesia was monitored frequently by testing the loss of the withdrawal reflex to a paw pinch. Eyes were moisturized using ophthalmic gel (GenTeal gel, Alcon), the analgesic meloxicam (Metacam, 5 mg/kg s.c.) was injected subcutaneously and the hair on the top of the head removed. Skin covering the skull was locally anaesthetized (5% lidocaine/prilocaine cream, Emla, AstraZeneca or bupivacaine 6-8 mg/kg s.c.), an incision made along the mid-line and the skill retracted to access the skull, which was cleaned using saline and absorbent triangles (Sugi). A craniotomy was then performed above the target brain region after identifying Bregma (SC: -3.4 mm posterior to Bregma, 0.5-0.8 mm lateral to Bregma; V1: -3.75 mm posterior to Bregma, 3 mm lateral to Bregma). Glass pipettes (Drummond) with a tip diameter of ∼20 μm were used to pressure-inject (Nanoject II; Drummond) viral suspension. Pipettes were backfilled with mineral oil (Sigma-Aldrich) and front-loaded with viral suspension. The pipette was lowered into the brain and viral injections were made at the following depths relative to the brain surface (SC: 2.0, 1.75, 1.5 and 1.25 mm, 220-280 nL total; V1: 0.8, 0.7 and 0.6 mm, 150 nL total) at a flow of 46 nL/s, with 1 minute between injections. After waiting at least 10 minutes following the last injection, the pipette was slowly retracted. The skin covering the skull was sutured (Ethilon Polyamide 6, Ethicon) and disinfected by topically applying a drop of antiseptic solution (Betadine). Mice were kept warm on a heating pad until fully recovered and monitored closely post-surgery. At least three weeks were allowed for viral expression before experiments were conducted.

### Intravitreal viral injections

Mice (male/female, 3-5 weeks of age) were anesthetized with isoflurane (3.5% induction; 1.8-2.0% maintenance in 0.3 L/min O_2_), placed on a feedback-controlled heating pad set to 37 °C (Harvard Instruments) and mounted in a stereotaxic apparatus (Kopf). The left eye ball of the mouse was carefully lifted out of its socket until the limbal plexus line could be observed and kept in place using a knotted thread around the eye. The pupil was dilated using 1-2 drops of atropine sulphate (Bausch and Lomb) to the surface of the eye. Once the pupil was fully dilated, a small hole was made at the dorso-temporal region of the eye using a 33G needle. This needle was then inserted 1/3 of the length of the bevel and 1 µL of vitreous fluid aspirated. A blunt 34G needle connected to a syringe (NanoFil, World Precision Instruments) was used to slowly inject 1 µL of virus suspension (AAV2-Syn-Chronos-GFP) mixed with a grain of methylene blue into the eye. Addition of methylene blue aided visualization of the viral suspension during eye injections. Ophthalmic gel (GenTeal gel, Alcon) was then applied to the eye, the thread removed and the eye was gently positioned back in the socket. Mice were kept warm on a heating pad until fully recovered and monitored closely post-surgery. At least three weeks were allowed for viral expression before experiments were conducted.

### Anesthetized animal preparation for *in vivo* electrophysiology

Mice (male/female, 4-12 weeks of age) were prepared for *in vivo* whole-cell electrophysiology as described previously^70^. Mice were anesthetized with a mixture of urethane (0.5-1 g/kg) and chlorprothixene (5 mg/kg) in saline by intraperitoneal injection. Depth of anesthesia was assessed using the paw withdrawal reflex. Mice were placed on a feedback-controlled heating pad set to 37 °C (Harvard Apparatus). Hair on top of the head was removed and the skin locally anesthetized (5% lidocaine/prilocaine cream, Emla, AstraZeneca). An incision was made along the midline, the skin over the skull was retracted and the skull was cleaned using saline. The skull was fixed to a custom-made head plate using UV-curing primer (Optibond All-In-One, Kerr) and dental cement (PermaFlo A1, Ultradent Products Inc.). A drop of silicon oil (Sigma-Aldrich) was applied to the eyes to keep them from drying out. Uni-or bilateral square craniotomies were made above the mediorostral region of the superior colliculus (0.8 mm rostral to Lambda [-3.4 mm posterior to Bregma], centered 0.5 mm lateral) using a dental drill (Osada Success 40, Osada). The dura was carefully removed using a 30G needle and forceps (Dumont type #7-curved), after which the brain surface was kept moist by application of artificial cerebrospinal fluid^71^ (aCSF), consisting of (in mM): 135 NaCl, 5.4 KCl, 5 HEPES, 1 MgCl_2_, 1.8 CaCl_2_ (pH 7.3).

### *In vivo* electrophysiological recordings

Electrophysiological recordings were performed using patch pipettes with a tip diameter of ∼1 μm and resistance of 4-8 MΩ. Pipettes were pulled from borosilicate glass (1.5 mm OD, 0.86 mm ID, Harvard Apparatus) using a P-97 micropipette puller (Sutter Instrument, USA) and mounted on a vertically-positioned headstage of a voltage and current clamp amplifier (Multiclamp 700A, Axon Instruments, USA). The amplifier headstage was attached to a remotely controlled micromanipulator (MP-285, Sutter Instrument). Recordings were digitized at 20 kHz (ITC-18, Instrutech, USA) and acquired using AxoGraph X software (v1.3.5, Axograph Scientific, Australia) running on an Apple Macintosh computer (iMac).

For juxtasomal recordings from SC neurons, recording pipettes were backfilled filled with aCSF and lowered to a depth of 1.0 mm below the pia under high pressure (>200 mbar), after which the pressure was decreased to 18-22 mbar. Recording pipettes were then advanced in 2 μm steps (2 μm/s) while monitoring the voltage trace in current clamp mode for spikes. For juxtasomal recordings from neurons in V1, recording pipettes were lowered to 0.5 mm below the pia under high pressure (>200 mbar), after which pressure was decreased to 18-22 mbar and the pipette advanced in 2 μm steps. Recordings were made from single-units with clear spike waveforms (>1 mV amplitude). For some experiments we used visual or optogenetic stimulation while advancing the pipette to increase the chance of finding single units. Spontaneous spiking was measured for >1 min before visual and optogenetic stimulation protocols were run.

*In vivo* whole-cell recordings were performed as previously described^70^. In short, patch pipettes were backfilled with intracellular solution, consisting of (in mM): 10 KCl, 130 K-gluconate, 10 HEPES, 4 Mg-ATP, 0.3 Na_2_-GTP, 15 Na_2_-phosphocreatine (pH 7.25-7.35, 290-300 mOsm). For recording from SC neurons pipettes were advanced to a depth of 1.0 mm below the pia under high pressure (>200 mbar), after which the pressure was decreased to 18-22 mbar. Pipettes were then advanced in 2 µm steps (2 µm/s) while monitoring the current response to 10 mV voltage pulses (at 50 Hz) in voltage-clamp mode. Upon a sudden increase in series resistance (detected as a sudden increase in the amplitude of the current response to the voltage pulse) positive pressure was removed and a gigaseal established (>1 GΩ resistance, -70 mV command voltage). After compensating for fast and slow capacitance transients, the whole-cell configuration was established by applying short suction pulses combined with the ‘Zap’ function (50-100 µs) of the Multiclamp amplifier. After applying bridge balance and capacitance neutralization in current-clamp mode we recorded the voltage response to current steps (-200 pA to +600 pA, 500 ms duration) before running visual and optogenetic stimulation protocols.

Electrophysiological parameters were calculated as follows. Membrane resistance and membrane capacitance were determined using the test-pulse protocol in AxoGraph. Series resistance represents the bridge balance value. Resting membrane potential was determined immediately after switching to current-clamp mode. Input resistance was determined based on the response to the smallest negative current step. Sag ratio was defined as the ratio between the peak negative response and the steady-state voltage at last 100 ms of the -200 pA current step. Rheobase was defined as the minimal current necessary to evoke action potentials. The following action potential parameters were determined based on the first 4-5 action potentials from near-rheobase current steps. Threshold was defined as the potential at which the action potential slope reached 20 mV/ms. After-hyperpolarisation amplitude was defined as the difference between the resting membrane potential and the peak negative potential after the downward sloping phase of the action potential. Action potential half-width was calculated at 50% of peak amplitude. Action potential rise-time was calculated between 20-80% of peak amplitude. Maximum action potential frequency was calculated from current injections before action potential failure occurred. At the end of *in vivo* experiments mice received a lethal dose of pentobarbitone solution (150 mg/kg i.p.), after which they were transcardially perfused with 4% paraformaldehyde (PFA) in 0.05 M phosphate-buffered saline (PBS). Brains were fixed overnight and stored in 30% sucrose at 4 or -20 °C.

### *In vitro* electrophysiological recordings

Acute brain slices were prepared from the SC in both hemispheres of adult mice (male/female, 6-7 weeks of age) at least 3 weeks after intravitreal injection of AAV2-Syn-Chronos-GFP into one eye. Mice were deeply anesthetized with isoflurane (3% in O_2_; 1 L/min). After testing for the loss of the withdrawal reflex to a paw pinch, mice were swiftly decapitated. The brain was extracted and sliced in ice-cold solution, containing (in mM): 110 Choline chloride, 11.6 Na-ascorbate, 7 MgCl_2_, 3.1 Na-pyruvate, 2.5 KCl, 1.2 NaH_2_PO_4_ and 0.5 CaCl_2_ bubbled with 95% O_2_/5% CO_2_ (pH of 7.4) using a vibratome (VT1200S, Leica Biosystems). Coronal slices (280 µm thick) were made and transferred to a holding chamber for 30 minutes at 35 °C in a solution containing (in mM): 92 NaCl, 2.5 KCl, 1.2 NaH_2_PO_4_, 30 NaHCO_3_, 20 HEPES, 25 glucose, 3 sodium pyruvate, 2 MgSO_4_ and 2 CaCl_2_ bubbled with 95% O_2_/5% CO_2_ (pH 7.4 ). Slices were then kept at room temperature. During recording slices were transferred to a recording chamber on an upright microscope (BX51; Olympus, Germany), held in place using a nylon-strung platinum harp, and perfused with aCSF at 30-32 °C, containing (in mM): 125 NaCl, 3 KCl, 1.25 NaH_2_PO_4_, 25 NaHCO_3_, 25 glucose, 2 CaCl_2_, and 1 MgCl_2_ bubbled with 95% O_2_/5% CO_2_ (pH of 7.4). Temperature was maintained using a feedback-controlled inline heater (Warner Instrument Corporation; TC-324B and SH-27B, United states). Cells were visualized using infrared differential interference (IR-DIC) optics and an infrared-sensitive camera^72^. Chronos-GFP expression in RGC axons was confirmed under 470 nm fluorescent light.

Whole-cell recordings were made from neurons in the SC, across multiple layers and depths. Patch pipettes with a tip diameter of ∼1 μm, a resistance of 5-6 MΩ and were backfilled with intracellular solution, containing (in mM): 135 K-gluconate, 7 NaCl, 10 HEPES, 2 MgCl_2_ and 2 mM Na_2-_ATP (pH 7.2, 290-300 mOsm). Pipettes were placed in a pipette holder connected to the head-stage of a patch-clamp amplifier (Dagan; BVC-700A, USA). Pipette position was controlled by a motorized micromanipulator (Luigs & Neuman; D-40880, Germany). Signals were digitized using an AD convertor (Instrutech; ITC-16 USA) and acquired on a computer (Apple iMac, USA) running AxoGraph X software (AxoGraph Scientific). AxoGraph X was used for both data acquisition and analysis. After establishing the whole-cell configuration, a current-step protocol (-200 to +100 pA, 20 pA steps) was performed for determining the active and passive electrophysiological properties of neurons. Postsynaptic responses to optogenetic stimulation were recorded in current-clamp mode. To block local network activity and thereby isolate monosynaptic responses slices were bathed in tetrodotoxin (TTX; 1 µM) and 4-Aminopyridine (4-AP; 100 µM).

### Visual stimuli

Full retina illumination was delivered using green LEDs placed over each eye (530 nm, ∼80 lux). The LEDs were shielded so that light was restricted to each eye separately. LEDs were connected to the analogue output of the digitizer (ITC-18, Instrutech, USA) and light intensity controlled by AxoGraph X software. LED stimulation protocols consisted of 25-75 trials with a trial duration of 4 seconds. On each trial, after a delay of 1 second to obtain baseline each eye was stimulated separately or together using a 20 ms LED light flash generated on a continuously background of low intensity illumination. Continuous low intensity, background illumination was used to mask potential ambient light and to prevent retinal activation during optogenetic stimulation.

Looming visual stimuli used during *in vivo* recording were generated in MATLAB (R2015b, MathWorks) using Psychtoolbox-3 on a dedicated high-performance Windows-based computer. Looming stimuli were identical to those used to trigger innate defensive behaviors, in that the stimulus was a 1 cm (2 visual degrees) black circle that rapidly widened to 25.5 cm (50 visual degrees) over 250 ms, after which it remained stationary for another 500 ms. Stimuli were presented to each eye individually or together using a haploscope consisting of two mirrors positioned at a 90° angle placed in front of the mouse. These mirrors reflected the image from two identical LED monitors (60 Hz frame rate, P2211H, Dell) placed on either side of head. Electrophysiological recordings were synchronized to visual stimuli by recording monitor pixel changes with a photodiode (BPW21R, Silicon PN, Vishay Semiconductors) located in the lower corner of the screen using AxoGraph X.

### Optogenetics

For optogenetic illumination of SC neurons *in vivo*, we placed an optic fiber (diameter 200 μm, 0.35 NA, 8 mm, Thorlabs) above the SC (-3.4 mm posterior to Bregma and 0.5 mm lateral) at a depth of 1 mm from the pia, corresponding to the dorsal surface of the SC. The optic fiber was connected to a LED light source (470 nm [M470F3] or 590 nm [M590F3] Thorlabs) and LED driver (LEDD1B, Thorlabs). Light was controlled by TTL pulses generated by AxoGraph X software. For optogenetic illumination of V1 neurons *in vivo*, the optic fiber was placed on top of the brain above V1 (-3.75 mm posterior to Bregma and 3 mm lateral). To prevent optogenetic light from activating the retina we optimized the strength of LED output (limited at 1.5 mW) and used low-intensity, background retinal illumination, as described above. Stimulus protocols consisted of 25-50 alternating visual trials with or without optogenetic stimulation. For optogenetic activation of RGC axons *in vitro* we used full-field illumination with a 470 nm LED (ThorLabs) mounted on the epi-fluorescent port of a microscope. Optogenetic activation of Chronos-expressing axons was done using 2 ms light pulses (intensity ranging between 0.5 mW and 5 mW).

### Data analysis of *in vivo* juxtasomal recordings

*In vivo* juxtasomal recordings were analyzed using custom-written MATLAB scripts. Only quality recordings (> 1 mV spike amplitude) with sufficient trials (> 25 trials) were included in the analysis. AxoGraph files were loaded into MATLAB using the ‘importaxo’ function (M.J. Russo, Kavli Institute, Columbia University, NY, USA). Raw traces were filtered using a Gaussian filter with a 500 Hz cut-off frequency (B.H.W. Winkelman, Netherlands Institute for Neuroscience, Amsterdam, NL) and spike timing was determined as negative or positive maxima of the first derivative of the voltage trace using the ‘findpeaks’ function. A manual quality-check was performed for each neuron to ensure correct spike timing. Continuous spike rates with 1 ms temporal resolution were calculated using a kernel density estimation (KDE) with a 10 ms gaussian kernel (N. Flierman, Netherlands Institute for Neuroscience, Amsterdam, NL). Spontaneous spike characteristics were calculated based on spike timing during spontaneous activity, or during the 1 s baseline period before visual stimulation (resulting in >50 s per neuron). Spike responses to visual stimulation were calculated by first normalizing KDE traces to the average spike rate 0.5 s prior to visual stimulation on a trial-by-trial basis. Maximum/peak spike rates were defined as the maximum of the KDE trace within a 0.5 s window from stimulus start. Latency to first spike was calculated as the median of trial-by-trial latencies between start stimulus and first spike timing (for trials in which spikes occurred within 0.5 s after start stimulus). Ocular dominance index (ODI) calculations were performed as described previously^33^.

### Data analysis *in vivo* whole-cell recordings

*In vivo* whole-cell data was processed first in AxoGraph and then analyzed using custom-written MATLAB scripts as follows. Neurons were analyzed if they had a stable resting membrane potential (-55.8 ± 1.3 mV) and spike amplitudes (57.3 ± 16.1 mV). In AxoGraph, individual trials were screened and excluded in case of >0.5 mV voltage fluctuations in the baseline before stimulus onset. To quantify subthreshold synaptic responses, traces were filtered using a 5 ms median filter, followed by a 1 kHz low-pass filter, after which the baseline during the 175 ms just prior to visual stimulation was subtracted. Files were then imported into MATLAB using the ‘importaxo’ function (M.J. Russo, Kavli Institute, Columbia University, NY, USA). The amplitude of EPSPs and IPSPs was determined during the 25-200 ms time period following stimulus onset. The area under synaptic responses was determined by calculating the sum of the absolute values during the same time period. EPSP/IPSP onset was calculated as the time between stimulus onset and 5% of response amplitude. To determine the time when contralateral synaptic responses differed from binocular responses, for each neuron we performed a cluster-corrected permutation test with 2000 permutations in MATLAB (E. Spaak, Oxford University, Oxford, 2015). To determine whether neurons showed significant negative or positive synaptic responses to visual or optogenetic stimulation we performed receiver operating characteristic (ROC) analysis on response amplitude values using custom code (S. Gharaei, Australian National University, Canberra). Heat plots were constructed using the MATLAB ‘imagesc’ function using a matrix sorted by the amplitude of responses as input. Cross-correlations were calculated using the MATLAB ‘xcorr’ function.

### K-means clustering and principal component analysis

K-means clustering and principal component analysis (PCA) on data acquired during *in vivo* whole-cell recordings was performed on 13 parameters: membrane resistance, membrane capacitance, resting membrane potential, input resistance, sag ratio, spike threshold, spike amplitude, spike rise time, spike half-width, spike after-hyperpolarization, maximum spike frequency, series resistance and recording depth. In some cells, missing values were substituted with the mean value obtained from the parameter dataset. K-means clustering was performed on the Z-score of superficially located SC neurons (depth 0-600 µm from the dorsal top of the SC) using the ‘kmeans’ function in MATLAB using 10,000 replicates. PCA was performed using the MATLAB ‘pca’ function.

### Data analysis *in vitro* whole-cell recordings

*In vitro* whole-cell data was processed in AxoGraph. Neurons showing resting membrane potential changes larger than 5 mV after wash-in of TTX/4-AP were excluded. Data from at least 10 repetitions were averaged and the baseline 0-20 ms before optogenetic stimulation subtracted. The amplitude of synaptic responses was determined between 20 and 100 ms after optogenetic stimulation. Neurons with EPSPs greater than 1 mV in response to optogenetic activation in the presence of TTX and 4-AP were classified as receiving monosynaptic RGC input.

### Histology and confocal imaging

Brains stored in 30% sucrose were frozen in embedding medium (Tissue-Tek O.C.T. compound, Sakura) and cut coronally at 50 μm or 100 μm thickness using a cryostat (Leica). Slices were washed in PBS, mounted on glass slides (Westlab) and covered using fluorescence mounting medium (DAKO) or fluorescence medium with DAPI (ProLong Diamond, Invitrogen). Images were acquired using a confocal microscope (SP5, Leica). For quantification of fluorescent of commissural axons (Figure 5), tilescans were performed with a 20x objective (Leica HC PL Apo, 0.75 NA), at 512 x 512 pixel resolution, 400 Hz scanning frequency with 2x line averaging. Laser power (514 nm for EYFP) and gain were kept identical between mice/slices. Subsequent quantification was performed using the Fiji distribution of ImageJ (v2.0.0-rc-69/1.52p). Fluorescence was measured within a 100 μm wide, vertical cylinder positioned 600-700 μm from the midline and averaged across the horizontal plane, resulting in a fluorescence-over-depth measurement (Figure 5C).

### Analysis of Allen Brain datasets

We quantified RGC axons fluorescence in images downloaded from the Allen Mouse Connectivity Atlas (connectivity.brain-map.org/projection) from the Allen Brain Institute^73^ via their application programming interface (API). The specific datasets used are: 306561747 (Cart-Tg1-Cre)^74^, 306957248 (Thy1-Cre), 306958034 (Thy1-Cre), 306930168 (Thy1-Cre) and 310035922 (Jam2-Cre). These five datasets are from mice (postnatal day (P) 91 to P102) injected intravitreally with enhanced green fluorescent protein (EGFP)-expressing adeno-associated viral vectors, with injection volumes ranging between 0.034-0.124 μL^75^. Projection density maps were acquired using two-photon serial tomography with 10 µm voxel resolution. Downloaded .nrrd data files were loaded into MATLAB and matched with the Allen Mouse Brain Common Coordinate Framework (CCFv3) 3D reference space with 10 µm voxel resolution. Slices within each dataset containing the SC (slice# 850-1000) were manually aligned (per 10 slices) to the reference images. Projection data was averaged across all datasets and heat plots representing expression in the anteroposterior and mediolateral directions were generated using the ‘imagesc’ function in MATLAB, after applying a 2-D Gaussian smoothing kernel with standard deviation of 0.5. To quantify expression in the dorsoventral axis, we first manually traced the dorsal border of the SC (per 5 slices) and used these tracing results to flatten the SC in the mediolateral axis in order to acquire accurate depth measurements. Quantification of fluorescence was done per hemisphere per individual mouse for slices containing the SC, by column-wise averaging non-zero values (expression in mediolateral axis), row-wise averaging non-zero values on the flattened SC (dorsoventral axis), and row and column averaging non-zero values (anteroposterior axis).

### Statistical analysis

Statistical tests were performed in SPSS (v24, IBM), Prism (v8.3.1 and v9.4.1, GraphPad) and MATLAB (R2022a, MathWorks). Normality was assessed using the Shapiro-Wilk test. Most data represented non-normal distributions, requiring non-parametric statistical tests (α = 0.05) as indicated for each test throughout the manuscript. Data are reported and presented as mean ± standard error of the mean (SEM), unless indicated otherwise. Differences were considered significant at *P < 0.05, **P <0.01 and***P < 0.001.

## Supplemental figures

**Figure S1.**
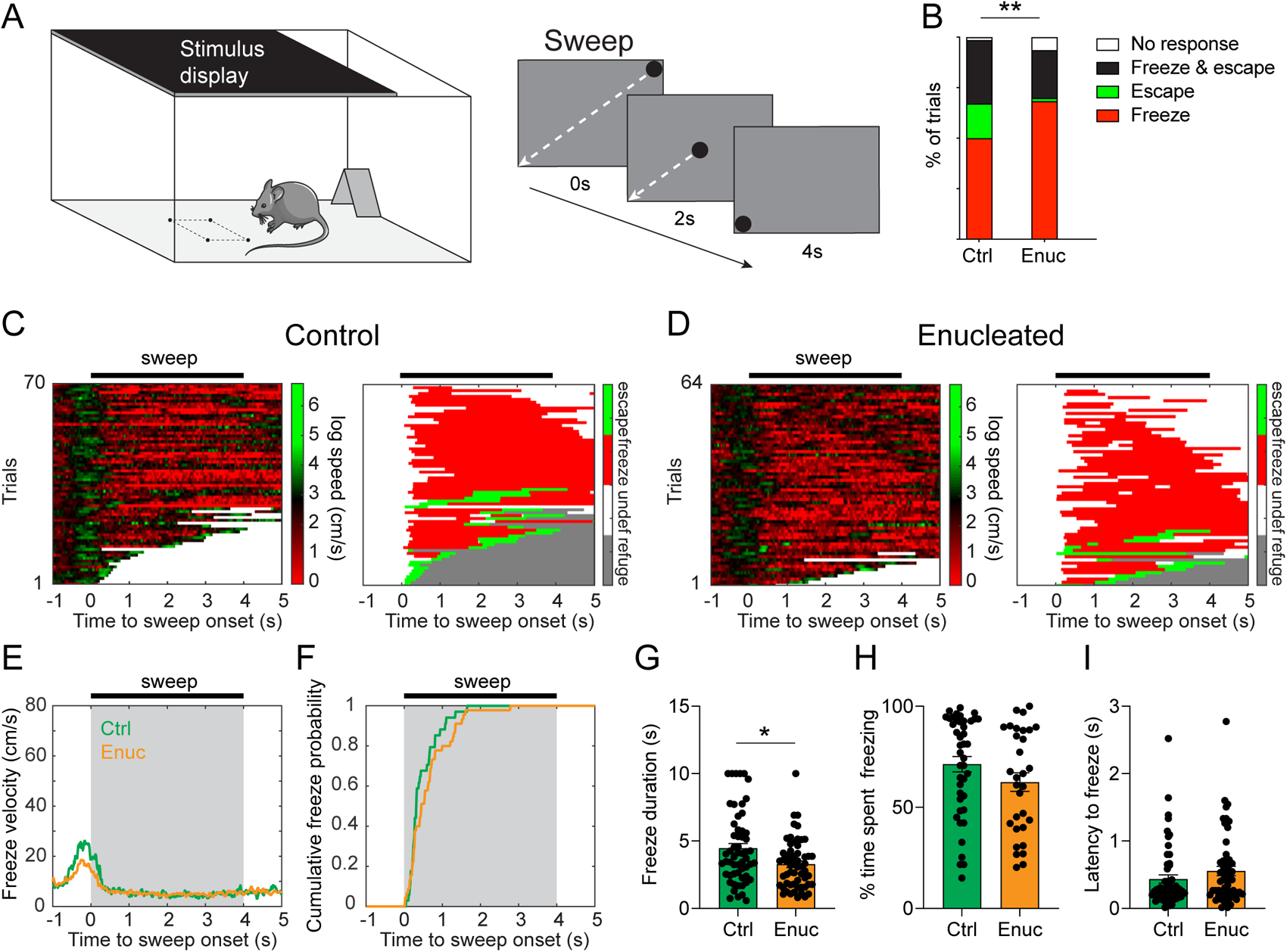
Binocular vision facilitates defensive responses to a moving spot. (A) Experimental setup for measuring defensive responses to visual stimuli. A single overhead moving spot (sweep), which moves from one diagonal corner of the screen to the opposite corner in 4s, was started when mice enter the center of the arena, indicated by the four black dots on the floor. (B) Responses of control (Ctrl) and enucleated (Enuc) mice across all trials. Control and enucleated mice showed primarily freeze responses, but control mice showed more escape, or freeze and escape, responses compared to enucleated mice (*χ* ^2^(3) = 12.92, p = 0.005, Chi-square test, n=70 trials from 15 Ctrl mice, n=63 trials from 17 Enuc mice). (C) Left: Heat map visualizing mouse movement speed relative to sweep onset across all trials of control mice. Right: Classification of responses relative to sweep onset. Undef (undefined) indicates time periods where behavior was not defined. (D) Same as in C, but for enucleated mice. (E) Median velocity on freeze trials relative to sweep onset. (F) Cumulative freeze probability relative to sweep onset. (G-I) Freeze duration (G; U = 1230, p = 0.018, Mann-Whitney *U* test, two-tailed, Ctrl: n=57 trials; Enuc: n=58 trials), percentage of time that mice spent freezing during sweep presentation (H; U = 521.5, p = 0.113, Mann-Whitney *U* test, two-tailed, Ctrl: n=43 trials; Enuc: n=31 trials) and latency (I; U = 1326, p = 0.161, Mann-Whitney *U* test, two-tailed, Ctrl: n=57 trials; Enuc: n=55 trials) on freeze trials. Data points show individual trials. Bar plots bars show mean ± SEM. *p < 0.05, **p < 0.01, ***p < 0.001.

**Figure S2.**
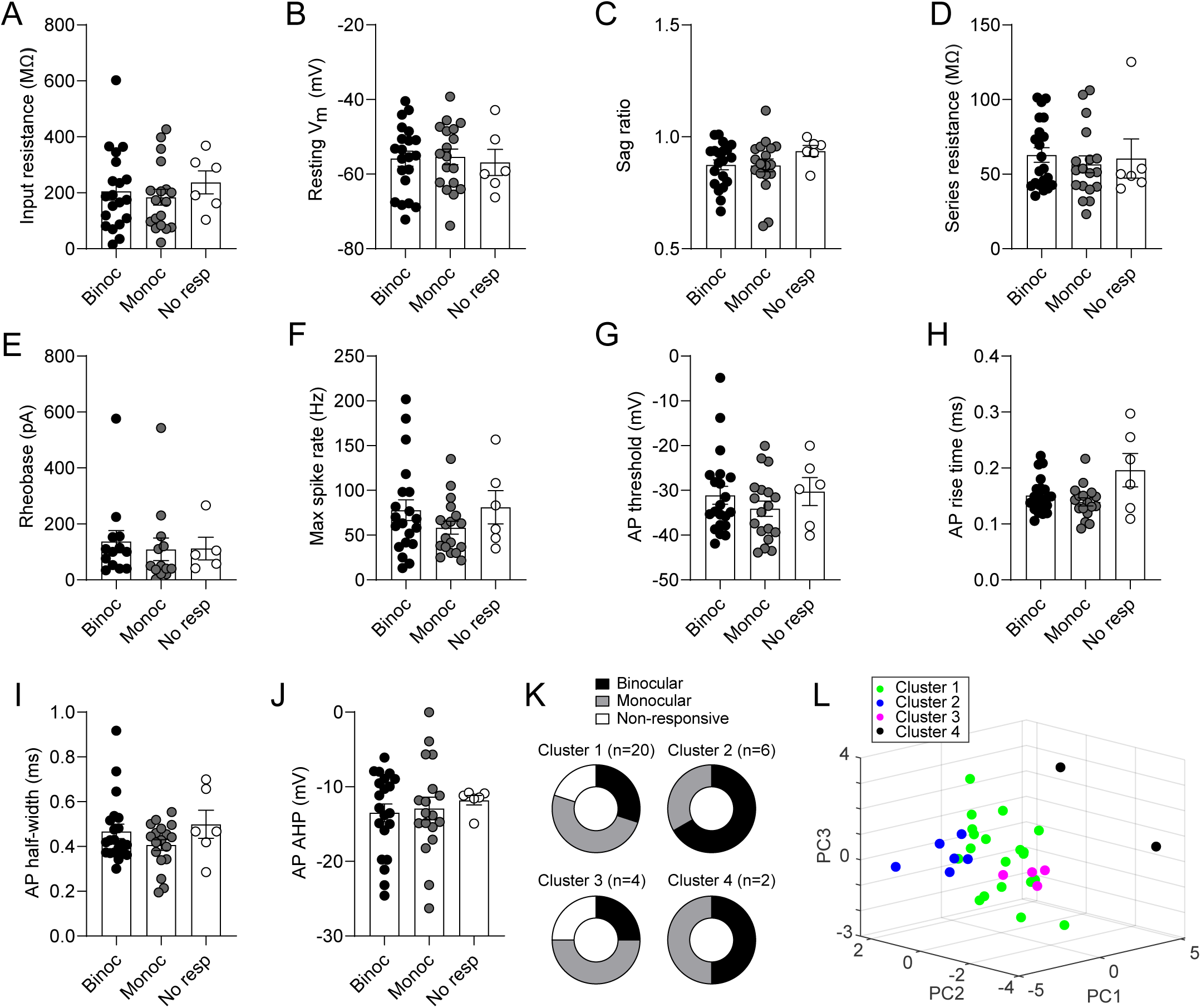
Electrophysiological properties of binocular, monocular and non-responsive neurons are similar. (A-J) Comparison of active and passive electrophysiological properties of binocular (Binoc), monocular (Monoc) and non-responsive (No resp) neurons. Statistical tests, corrected for multiple comparisons, show no significant difference for any of the parameters. V_m_ = membrane potential; AP = action potential; AHP = after-hyperpolarization; sag ratio = ratio between the negative peak and steady state during strong negative current injection. (K) Binocular neurons occur in every category determined by k-means clustering based on all 13 electrophysiological properties, indicating that binocular neurons are not limited to specific superficial superior colliculus cell-types. (L) Plot showing the first 3 principal components after performing principal component analysis (PCA). Colors indicate the 4 categories that cluster together in PCA1-3 parameter space. Data points show individual neurons. Bar plots show mean ± SEM.

**Figure S3.**
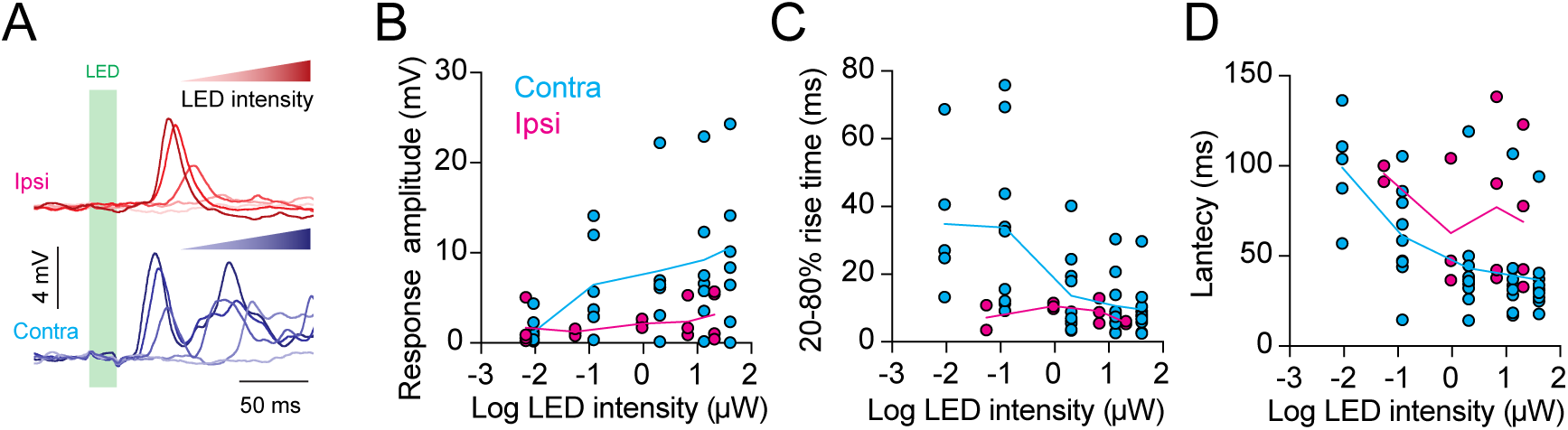
Binocular synaptic responses scale with LED intensity. (A) Example visually-evoked responses of a representative binocular neuron to increasing LED intensity applied separately to the contralateral and ipsilateral eye. (B-D) Amplitude (B), 20-80% rise time (C) and latency (D) of EPSPs in SC neurons evoked by LED contralateral and ipsilateral eye stimulation of varying intensity. Data points show neurons (n=3 ipsilateral, n=5 contralateral).

**Figure S4.**
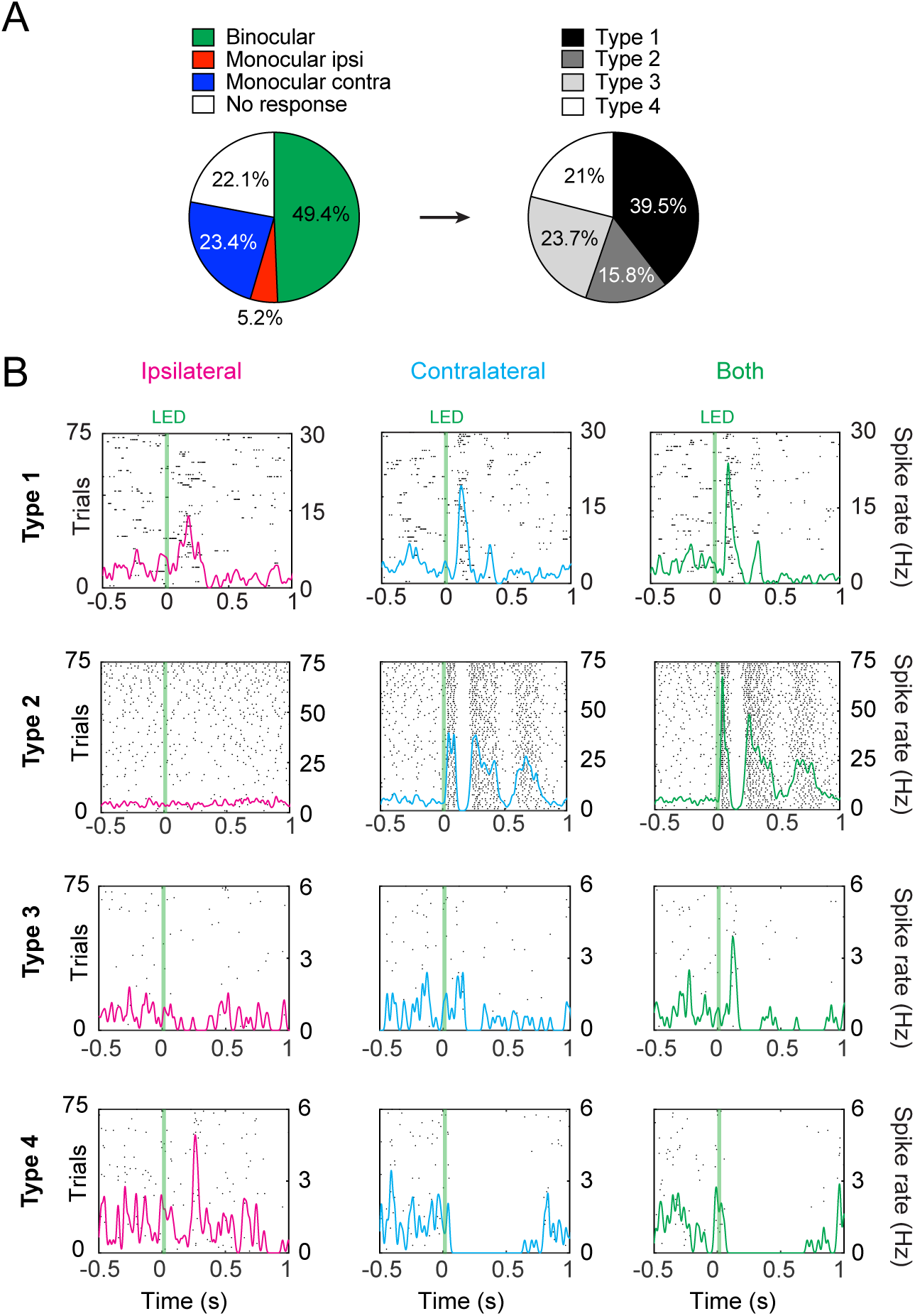
Binocular spiking types in superior colliculus. (A) Distribution of spike responses across all recorded neurons (left; n=77 neurons from 5 mice). Binocular neurons were further sub-divided into 4 different types based on their spiking response (right; see B). (B) Peristimulus time histograms of the 4 binocular spiking response types (1-4; rows) during stimulation of the ipsilateral or contralateral eye alone, or both eyes together (columns). Each dot representing a spike. Colored lines show spike rate (right Y-axis). Type 1 neurons had spiking responses to stimulation of the ipsilateral and contralateral eye on their own. Type 2 neurons had spiking responses to stimulation of only the ipsilateral or contralateral eye, but the response to stimulation of both eyes together was different to the response during stimulation of the responsive eye. Type 3 neurons do not have a spiking response to stimulation of either eye alone, but had a spiking response to stimulation of both eyes together. Type 4 neurons had a spiking response to stimulation of one eye, and inhibition of spiking during stimulation of the other eye, with inhibition of spiking during stimulation of both eyes together.

**Figure S5.**
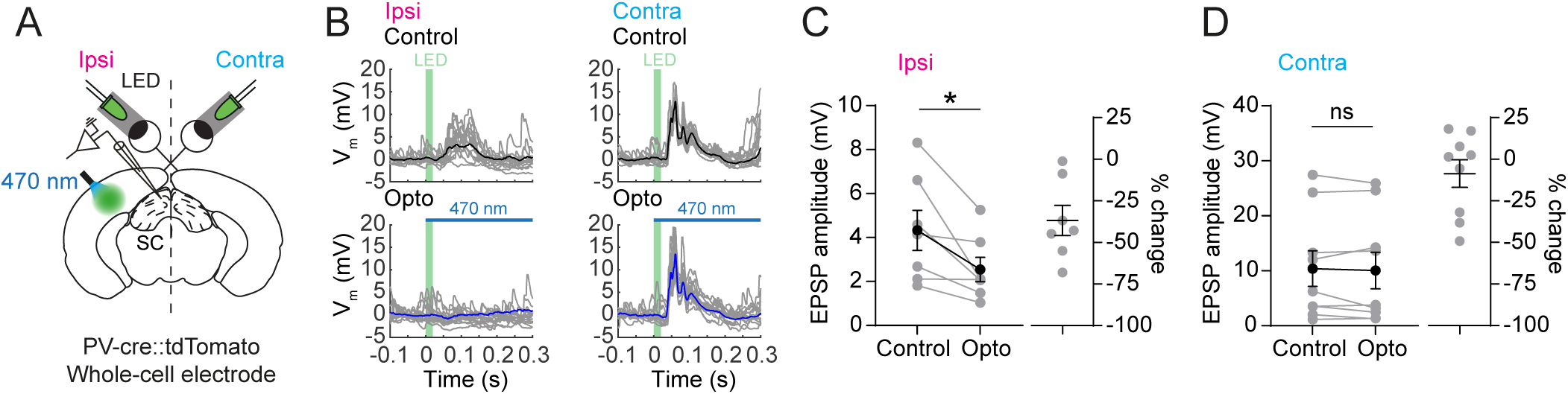
Silencing of primary visual cortex neurons reduces ipsilateral synaptic responses in superior colliculus. (A) Schematic of *in vivo* whole-cell recordings from the SC ipsilateral to the binocular V1 injected with cre-dependent ChR2 virus during LED stimulation of the eyes and optogenetic activation of inhibitory Parvalbumin (PV) neurons through an optic fiber emitting 470 nm light. (B) EPSPs during ipsi-(left) and contralateral (right) eye stimulation without (‘control’, top) and with (‘opto’,bottom) optogenetic silencing of binocular V1. (C) Optogenetic silencing of binocular V1 leads to a decrease in ipsilateral eye EPSPs (amplitude: -37 ± 9%, *t*(6) = 2.88, p = 0.028, area: -19 ± 5%, *t*(6) = 3.041, p = 0.023, paired *t*-test, two-tailed, n=7 neurons from 6 mice). (G) Contralateral eye EPSPs are not changed (amplitude: W = -9, p = 0.652; area: W = 19, p = 0.3, Wilcoxon matched-pairs signed rank test, two-tailed, n=9 neurons). Data points show individual neurons. Lines and error bars show mean ± SEM. *p < 0.05, **p < 0.01.

